# Are your data too coarse for speed estimation? Diffusion rates as an alternative measure of animal movement

**DOI:** 10.1101/2025.07.17.665364

**Authors:** Vickie L. DeNicola, Stefano Mezzini, Francesca Cagnacci, Christen H. Fleming

## Abstract

Estimates of speed and distance traveled are routine in ecological research to provide a link between behavior and energetics. Conventional straight-line displacement (SLD) methods return severely and differentially biased estimates with no ability to evaluate the accuracy of the estimate. Recent methodological advances have improved our ability to estimate these parameters using continuous-time speed and distance (CTSD) estimation. However, even with CTSD estimation, many datasets are too coarse, or the location error is too great to reliably measure speed or distance traveled. To address these limitations, we investigated the relationship between CTSD-estimated mean speed and diffusion rate, where diffusion rate is defined as the variance in displacements per time interval. We calculated CTSD mean speed and diffusion rate estimates using telemetry data (i.e., trajectory data) from over 100 white-tailed deer and simulated telemetry data generated from a known movement model. We examined the relationship between the two measures in both datasets and the effect of sampling frequency on the effective sample size and the estimation of the two parameters. We found that mean speed and diffusion rate were strongly and nonlinearly correlated, with a 1% increase in diffusion rate predicting a 0.40% increase in mean speed (99% CI: 0.38–0.42%) for our focal species. Diffusion rate estimates remained substantially more accurate and precise than speed across sampling interval regimes, even when speed estimation was not possible. Our findings demonstrate that diffusion rate outperforms mean speed as a measure of movement activity under marginal data conditions. Diffusion rate is a reliable measure of movement activity and can link behavior to energetics across a wider range of datasets while maintaining accuracy even when data quality is low. By using speed and diffusion together, researchers can rely on more robust insights into space use across a range of ecological contexts.

## 1. INTRODUCTION

Animal movement is a fundamental and rapid dynamic response to environmental or physiological stimuli (Nathan et al., 2008), driven by energetic requirements, reproductive needs, competitor and predator avoidance, habitat fragmentation, and other processes operating at various spatial and temporal scales (Carbone et al., 2005; Nathan et al., 2008; Schlägel et al., 2020; Van Moorter et al., 2016). Measures of how much and how fast animals move are central to optimal foraging theory (Brown et al., 1999) and the marginal value theorem (Charnov, 1976), as both frameworks emphasize the trade-off between energetic costs and resource acquisition.

Movement serves as the bridge between these costs and benefits, with distance and speed influencing the time and energy required to locate, access, and consume resources. The marginal value theorem predicts that animals should leave a foraging patch once resource intake falls below the environment’s average intake rate — a decision shaped by movement speed and distance. Similarly, optimal foraging theory suggests that animals adjust movement patterns to balance exploration for new resources with exploitation of known patches. As a result, estimates of distance traveled and speed are widely used in ecological research as proxies for the energetic costs of movement, providing a tangible link between theory, behavior, and energetic efficiency (Garland, 1983; Wilson et al., 2008).

Often, distance traveled is estimated using the linear distance between consecutive locations, referred to as straight-line displacement (SLD), and speed is estimated by dividing an SLD by the time elapsed between the corresponding pair of locations. This form of estimation is straightforward to calculate, but it results in biased estimates of both distance traveled and speed (Noonan et al., 2019; Rowcliffe et al., 2012; Sennhenn-Reulen et al., 2017). SLD-derived estimates fail to account for the inherently tortuous and non-linear nature of many animal trajectories (Nathan et al., 2022; Fig. 2a in Noonan et al., 2019), resulting in a substantial underestimation of the true distance traveled and speed (Noonan et al., 2019). This underestimation occurs because an animal’s movement trajectory is a latent variable: it is not possible to know exactly the true single steps building up the movement path. While one can approximate that using sequences of telemetry-based locations (e.g., GPS locations, Nathan et al. 2008; henceforth “telemetries”), such data only provide partial information on where and how the animal moved (Fleming et al., 2016). Thus, SLD underestimates speed and distance traveled, particularly when animals exhibit complex behaviors, such as foraging or social interactions, at a timescale that is finer than the sampling interval of the locations. This is exacerbated by coarse sampling and data thinning, which is regularly used to homogenize sampling interval (Rowcliffe et al., 2012). Conversely, fine-scale sampling mitigates this underestimation but can also result in an overestimation driven by telemetry measurement error (Fig. 2b Noonan et al., 2019; Rowcliffe et al., 2012). Finally, although the SLD method can be modified to account for these measurement error biases (Johnson et al., 2011), fine-scale sampling introduces autocorrelation structures that cannot be accounted for using SLD methods (Noonan et al., 2019). Additionally, applying conventional SLD methods will return speed and distance traveled estimates without confidence intervals or uncertainty estimates (Noonan et al., 2019).

Continuous-time speed and distance (CTSD) estimation is an alternative to SLD-based estimation that corrects the issues mentioned above. It is based on continuous-time stochastic movement models (Fleming et al., 2014; Gurarie et al., 2009) that leverage the autocorrelation in the data to inform both tortuosity and measurement error simultaneously (Fleming et al., 2014, 2016; Noonan et al., 2019). By accounting for the non-independence between locations, CTSD estimation produces estimates of effective sample size (ESS) for movement parameters, including speed, home-range crossing time, and home-range size. ESS is a measure of the number of independent samples for each parameter estimate and ensures that parameter estimates and confidence intervals reflect the true informational content of the data and avoid biases caused by unmodeled autocorrelation (Fleming & Calabrese, 2017). Recent applications of CTSD estimation have demonstrated its ability to accurately reconstruct movement behaviors and allow ecologists to understand behavioral processes at fine ecological timescales (Nathan et al., 2022; Noonan et al., 2019; Thompson et al., 2021). However, CTSD estimation only provides accurate estimates when the data are sufficiently fine and the location error is sufficiently small to inform the speed parameter estimates (Noonan et al., 2019).

This study introduces continuous-time estimated *diffusion rate* as a robust and versatile metric for quantifying animal movement and proposes it as an alternative to discrete and continuous time estimates of mean speed, especially for coarse telemetry data. We define diffusion rate as the expected square displacement over a finite period of time, which can be interpreted as the variance in locations per unit of time (Supplement A). Unlike speed, diffusion rates offer insights into how extensively an animal explores space over time (Moorcroft & Lewis, 2013, allowing researchers to investigate cumulative displacement patterns less sensitive to sampling bias.

This paper had two main aims: (1) to demonstrate and describe the relationship between mean speed and diffusion in animal trajectories and (2) to demonstrate why and when diffusion rate is a better metric of animal movement. Our analysis draws on empirical tracking data from a large dataset of 108 white-tailed deer (117 seasonal trajectories; *Odocoileus virginianus* Zimmermann; hereafter, deer) from New York, USA. We applied CTSD models to generate estimates of mean speed and diffusion rate and demonstrate the relationship between the two as well as how sampling frequency affects their estimates and effective sample size. Finally, to show how speed estimation performs under increasingly coarse conditions, we used simulations from a known movement model to assess the accuracy and robustness of both mean speed and diffusion rate as a function of sampling interval. This study demonstrates the broader applicability of diffusion rate in ecological research and provides a foundation for more reliable movement modeling.

## 2. METHODS

Diffusion rates measure how broadly an animal’s location spreads over time by quantifying the rate of increase in spatial variance with respect to time, providing insight into how quickly an animal expands its range (Okubo & Levin, 2001). The diffusion rate D can be defined as the variance in displacements per unit time-lag or

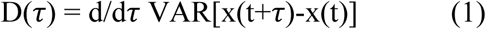

where *x(t)* is the location process at some initial time t, and τ is the time lag under consideration (Okubo & Levin, 2001). Unlike mean speed, which captures the average rate of movement per unit time, the diffusion rate considers both speed and directional changes. Mean speed or daily distance traveled can provide an important indicator of an animal’s energetic expenditure (Carbone et al., 2005), whereas diffusion rate can inform on space use and movement efficiency (Okubo & Levin, 2001).

### 2.1 Relationship between mean speed and diffusion rate in deer telemetries

To quantify the relationship between speed and diffusion, we used the mgcv package (Wood, 2017) for R (R Core Team, 2024) to fit a Hierarchical Generalized Additive Model (HGAM; Pedersen et al., 2019) to the moving-window estimates of mean speed and diffusion produced in DeNicola et al. (2025) (see Supplement B). To remove potential confounds due to differences in sampling intervals (discussed below), we constrained the analyses to windows with a median sampling interval of 1 hour (95.13% of the estimates with finite speed and diffusion). The HGAM had a population-level linear term of log-transformed diffusion and a factor smooth term of log-transformed diffusion for each animal to account for individual-level deviations from the population average (see model GS in Pedersen et al., 2019). The model had a Gamma family of distributions with a log link function, so the population-level model (i.e., ignoring individual animal-level trends) can be written as

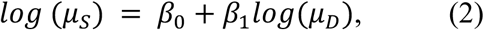

where μ_S_ is mean speed and μ_D_ is mean diffusion. We discretized the covariates (discrete = TRUE) and the smoothness parameter was estimated via fast REML (method = “fREML”) to reduce fitting time with no substantial loss to accuracy.

### 2.2 Effects of sampling interval on robustness of estimates in deer telemetries

To estimate the effects of sampling interval on the relationship between mean speed and diffusion as well as one’s ability to obtain estimates of speed and diffusion rate, we thinned all trajectories with progressively coarser sampling intervals. Starting from the estimates that had a median sampling interval of 1 hour (117 seasonal trajectories from 108 distinct deer, with 9 tracked in two separate seasons), we thinned the data at intervals of 2, 3, 4, …, 46, 47, and 48 hours and fit movement models to each trajectory using the ctmm.select() function for a total of 5,616 movement models. We then modeled the effects of diffusion and sampling interval on mean speed using a second HGAM. In addition to the terms from model (2), this second HGAM included a population-level smooth term of sampling interval. As with the first HGAM, this model used a Gamma family of distributions and a log link function, and the smoothing parameter was estimated via fast REML with discretization of the covariates. We can thus write the population-level model as

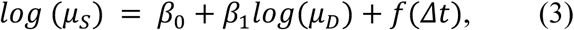

where *f*(Δt) is a (smooth) function of the sampling interval.

### 2.3 Relationship between speed and diffusion rate in simulated trajectories

To demonstrate the consequences of forcing speed estimation when data are too coarse, we simulated 1,000 trajectories from a known movement model. The model had a directional persistence of 1 hour, a range crossing time of 24 hours, and a spatial variance of 1 km^2^, which resulted in a mean speed of 6.14 km/h and a mean diffusion of 1.74 km^2^/h. We thinned each of the 1,000 trajectories as above using sampling intervals of 1, 2, 3, …, 46, 47, and 48 hours. We then fit Ornstein-Uhlenbeck-Foraging (OUF) movement models via the ctmm.fit() function to obtain estimates of both speed and diffusion rate for all sampling intervals, forcing estimates for coarse data.

## 3. RESULTS

The deer had a mean estimated directional persistence of 10.2 minutes (median 9.7 minutes; range: 2.5-26.9 minutes, all below the sampling interval of 1 hour) and a mean estimated range crossing time of 27.8 hours (median 9.1 hours; range: 3.7 hours to 33 days).

### 3.1 Relationship between mean speed and diffusion rate in deer telemetries

The model for the 1-hour moving window data demonstrated a good fit, explained 80.9% of the deviance, and did not show evidence of heteroskedasticity in the residuals (Fig. 1). It produced the estimated relationship in SI units

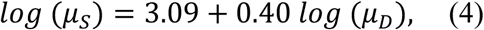

where μ_S_and μ_D_ are the estimated mean speed and diffusion in SI units, respectively. The equation can be rewritten as the estimated power relationship from

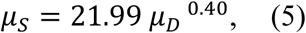

which we can derive that speed increases with diffusion sublinearly (since the 0.40 < 1), and a 1% increase in diffusion results in an approximate 0.40% increase in mean speed (99% CI: (0.38, 0.42); see Supplement C).

**Figure 1.**
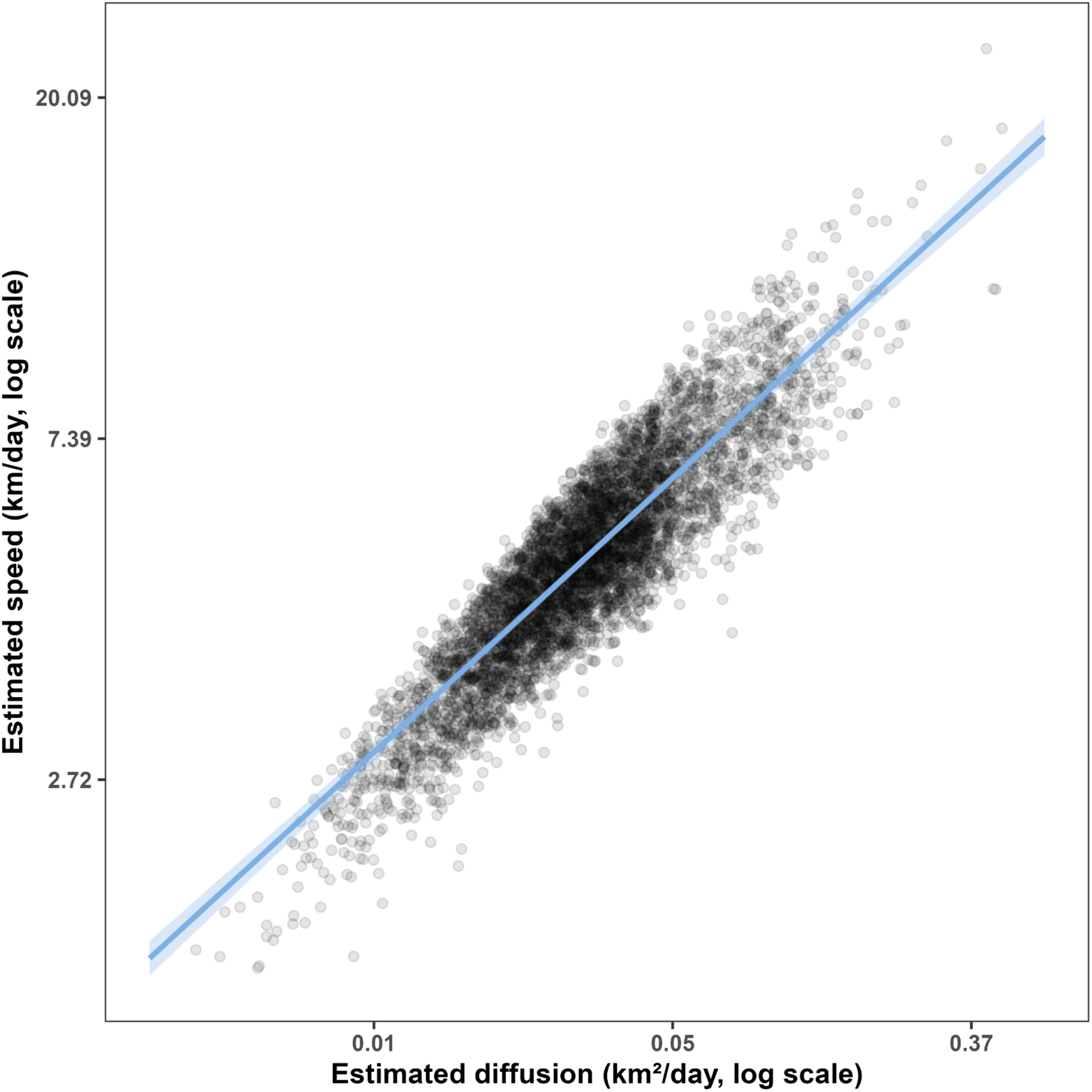
Estimated speed and diffusion have a linear relationship on a log-log scale for deer telemetries. Movement rates were estimated using continuous-time movement models using the ctmm.select() function from the ctmm package for R. The line indicates the estimated relationship, while the shaded ribbon indicates the 99% credible interval of the posterior. The points are 5,180 7-day mean speed and diffusion estimates from 108 deer estimated in DeNicola et al. (2025).

### 3.2 Effects of sampling interval on robustness of estimates in deer telemetries

Of the 5,616 movement models fit to the thinned deer telemetries, only 984 (17.5%) had finite estimates of speed, but 4219 (75.1%) had finite estimates of diffusion. All models with finite speed also had finite diffusion. The HGAM fit to these data revealed a similar relationship to the one described in Eq. (5) after accounting for the effect of sampling interval (Fig. 2). However, the relationship between mean speed, μ_!_, and sampling interval, Δt, was nonlinear, so we included its effect as the nonlinear (smooth) function *f*(Δt), which can be treated as a constant for a given Δt. The model produced the power relationship and

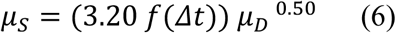

had a very good fit, as it explained 97.7% of the deviance. The estimated exponent was 0.504 (99% credible interval: (0.463, 0.545)).

**Figure 2.**
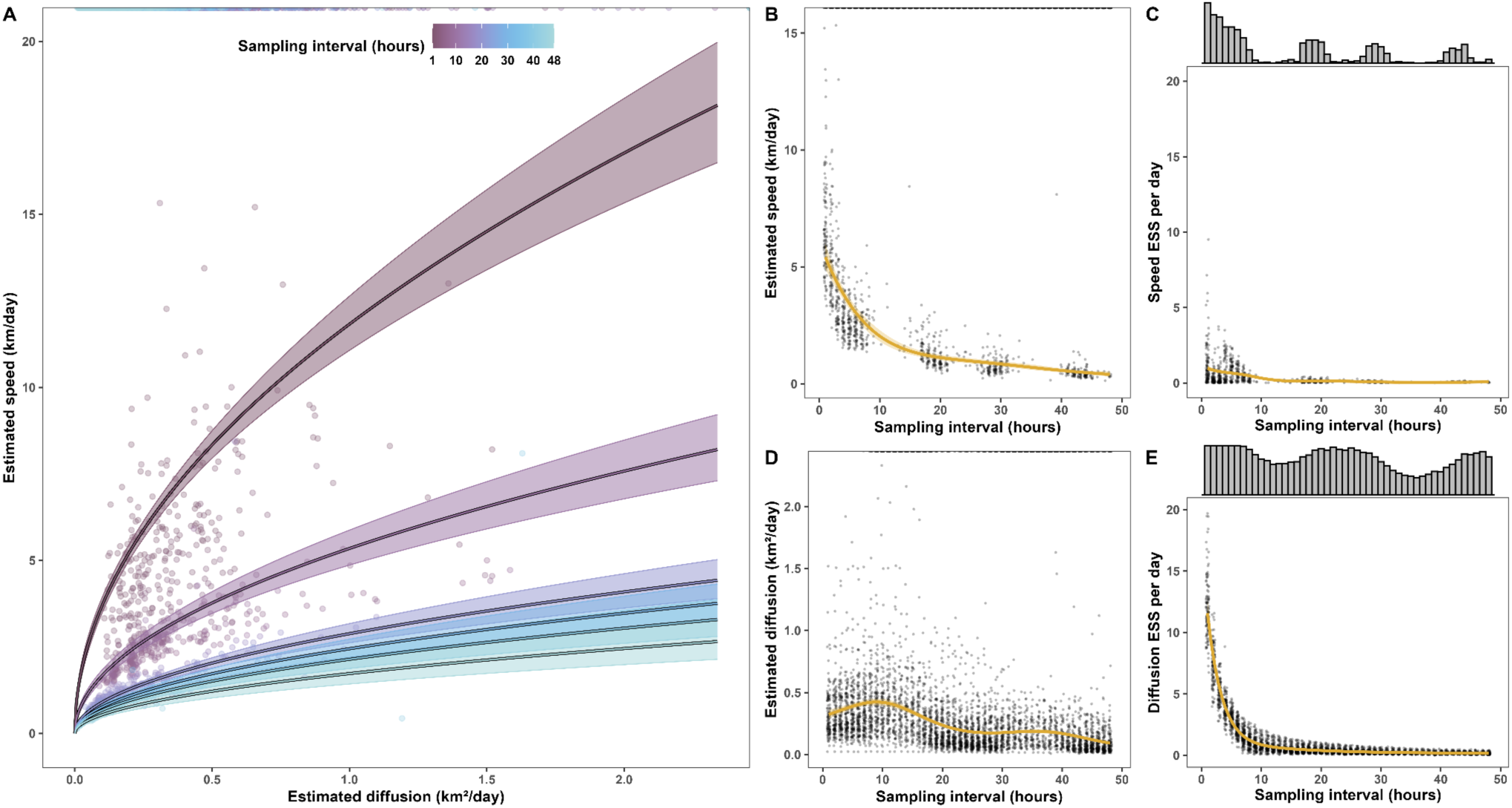
Estimated speed and diffusion estimated from deer telemetries have a nonlinear power relationship (A). The line indicates the estimated relationship, while the shaded ribbon indicates the 99% credible interval. The points at the top of the figure show the distribution of non-finite estimates of speed (top) and diffusion (top right corner). Speed estimates (B) and effective sample size (C) decrease rapidly with sampling interval and show substantial gaps where speed estimation was inappropriate (see marginal histograms in C). Diffusion estimates (D) and effective sample size (E) also decrease with sampling interval. Still, diffusion estimates do not change substantially for sampling intervals ≲ 15 hours, effective sample sizes are much higher than those for speed, and the estimate gaps are less pronounced. Orange lines and ribbons indicate the relationships estimated by Gamma GAMs fit via ggplot2’s geom_smooth() function, while points at the top of panels B and D indicate non-finite speed and diffusion estimates, respectively. Marginal histograms in C and E indicate the number of finite speed and diffusion estimates, respectively.

Diffusion estimates and effective sample size were substantially more robust to sampling interval than speed estimates. Mean-speed estimates decreased with sampling interval, and mean-speed effective sample size was > 5 in only 0.08% of the estimates. In contrast, diffusion rate estimates did not change substantially across sampling intervals ≲ 15 hours. Approximately 5.48% of all thinned telemetries had a diffusion effective sample size > 5, including all those with a 1-hour sampling interval. Both parameters showed oscillations in the number of models with finite estimates, but the oscillations were much more accentuated for mean speed. All sampling intervals had more than 40 diffusion estimates (out of 117 movement models), but 41 of 48 sampling intervals had only 40 speed estimates or less.

### 3.3 Relationship between speed and diffusion rate in simulated trajectories

The results from the simulated trajectories mirror those from the previous two subsections: Speed estimates were reasonably accurate and precise for sampling intervals ≲ 4 hours (Fig. 3A; also see Noonan et al., 2019), while diffusion estimates were reasonably accurate and precise for sampling intervals ≲ 23 hours (Fig. 3B). Sampling intervals as low as 6 hours resulted in infinite speed point estimates, while 99.82% of all diffusion-rate point estimates were finite, albeit often excessively large for sampling intervals ≥ 35 hours. As in the previous section, effective sample size for speed decreased faster over the sampling interval than for diffusion, but note that it was substantially larger when the sampling interval was near the true directional persistence timescale of 1 hour (Fig. 3C-D).

**Figure 3.**
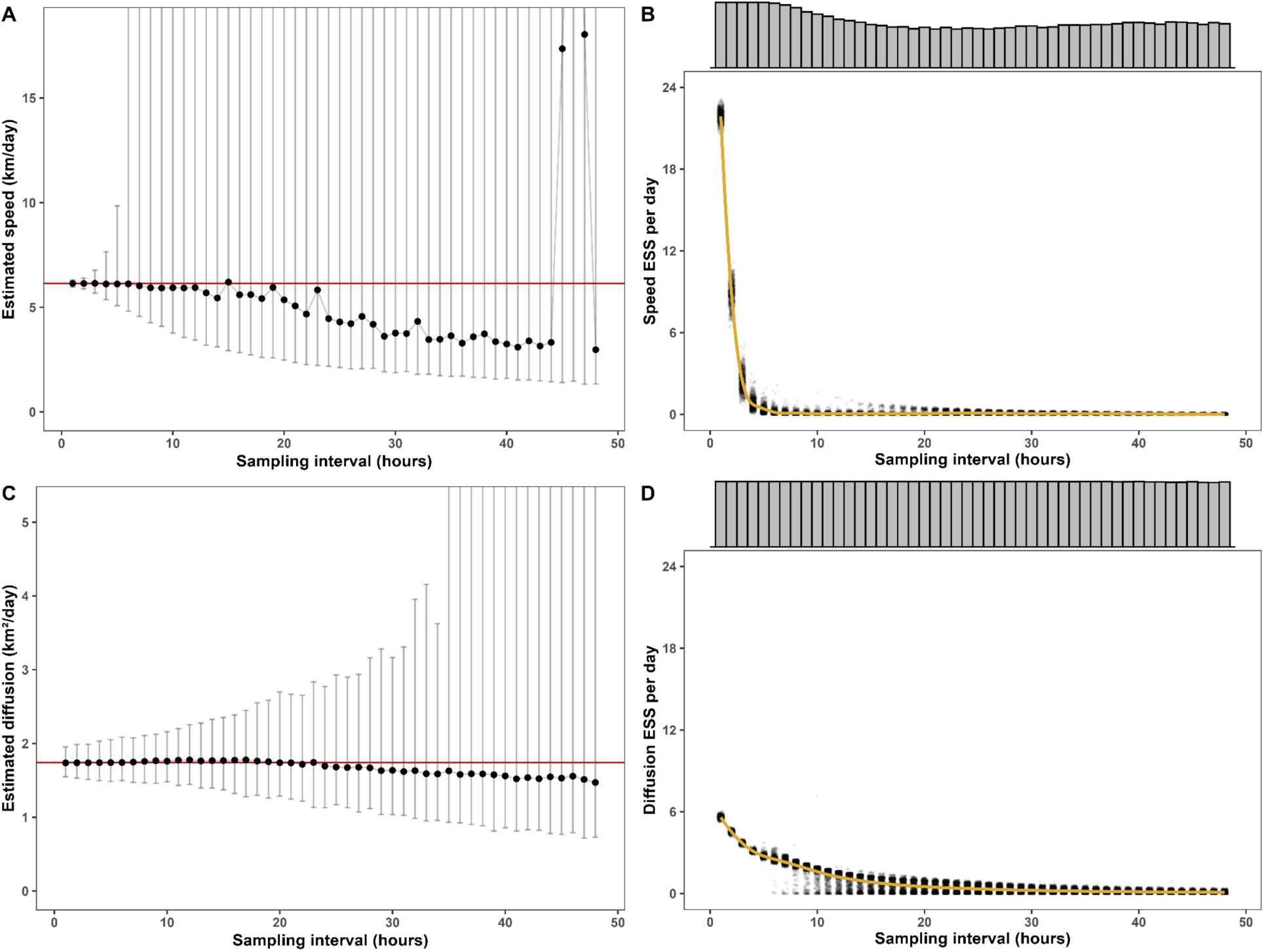
Estimated speed (A) and diffusion (C) and corresponding estimated effective sample sizes (ESS) for speed (B) and diffusion (D) from 1,000 Ornstein-Uhlenbeck-Foraging (OUF) movement models fit via the ctmm.fit() function from the ctmm package to simulated trajectories. The points and error bars in panels A and C indicate the median estimate and the 99% quantile interval, respectively, while the red line indicates the true speed and diffusion parameters. In panels B and D, the points indicate the data, jittered horizontally by no more than ±0.25, while the orange line indicates the estimated trends fit via ggplot2’s geom_smooth() function. The marginal histograms indicate the proportion of finite estimates at each sampling interval.

## 4. DISCUSSION

Fine-scale animal movement is often characterized using speed and distance traveled to answer questions about foraging patterns, resting habits, and travel routes while linking movement intensity to an animal’s responses to environmental stimuli (Fryxell et al., 2008; Turchin, 1998; Wilson et al., 2015). An animal’s diffusion rate reflects the individual’s short-term space use by integrating movement speed and tortuosity to reveal larger-scale processes such as site fidelity or exploration (Codling et al., 2008; Smouse et al., 2010). Higher diffusion rates suggest exploratory behavior or dispersal, whereas lower rates indicate either inactivity or more localized activities, such as the use of den sites, foraging patches, or breeding behavior (Codling et al., 2008; Giuggioli & Kenkre, 2014). Taken together, speed and diffusion rate provide complementary insights into an individual’s movement behavior. Speed captures the rate and intensity of movement, while diffusion links movement to broader spatial and ecological processes (Börger et al., 2008). Using both metrics together, researchers can effectively characterize the proximate behaviors driving movement and the ultimate ecological consequences of space use. Additionally, when accurate speed estimation is not possible due to limitations such as coarse sampling intervals, measurement errors, or low effective sample sizes, diffusion rate can be used as a robust proxy for mean speed. While diffusion rate does not directly measure speed, it incorporates information on movement rate and directional persistence, offering insights into overall movement activity. Our findings highlight that, at coarse sampling intervals, diffusion rate is a more accurate, robust, and consistent measure of an animal’s movement behavior that can be used in place of mean speed.

### 4.1 Relationship between mean speed and diffusion rate in deer telemetries

Our results showed that mean speed and diffusion rate scale with a sublinear power relationship, suggesting that diffusion can be used to make population-level inferences about speed. However, this relationship does depend on movement behavior at the individual level. Since diffusion reflects the combined effects of speed and tortuosity, it can provide a better proxy for mean speed when tortuosity is less variable in the population, whether that be an ecological population or a statistical population of different behaviors. Tortuosity can vary from predominantly ballistic movement (e.g., moving between patches without exploring much) with low tortuosity and high directional persistence to predominantly exploratory movement (e.g., moving within patches) with high tortuosity and low directional persistence. Animals exhibiting highly directed ballistic movement (Visser & Kiørboe, 2006) will generate both large diffusion rates and mean speeds. In contrast, animals with highly tortuous movement that explore a large area will have high mean speeds but comparatively lower diffusion rates, as the space they fill does not scale linearly with speed. Therefore, all else being equal, increasing tortuosity will increase mean speed without substantially impacting the diffusion rate. This relationship highlights how movement characteristics—such as directional persistence and tortuosity—shape the scaling of speed and diffusion rates.

We expected a positive association between mean speed and diffusion rate within populations, and our results suggest this association is approximately linear on a log-log scale (i.e., sublinear on natural scales). However, this relationship may vary depending on the extent to which mean speed and tortuosity differ across individuals within the population, given differences in movement behavior and physiology. Further research on a broader range of species is needed to determine whether diffusion rate remains a simple proxy for mean speed and distance traveled at the population level. While diffusion rate is a measure of large-scale movement patterns and a useful alternative when speed estimates are unavailable or unreliable, it cannot be assumed to be proportional to speed or used as a direct proxy without consideration of the nonlinear scaling between the two variables and species- or individual-specific movement characteristics.

### 4.2 Effects of sampling interval on robustness of estimates in deer telemetries

Our results demonstrated that diffusion rate is substantially more robust to coarse sampling intervals than speed and that it is easier to obtain realistic and accurate diffusion estimates under marginal data conditions. However, the oscillations in the mean and number of estimates over the sampling interval suggest that the ability to produce accurate estimates of diffusion (or any estimates at all) depends in part on the animals’ heterogeneity in movement behavior. Unlike the simulations we present in section 3.3 and discuss below, animals do not have stochastic movement behaviors with a single mean. Instead, they often switch between behaviors (i.e., many different means, e.g., foraging, exploration, sleeping), so estimates of any movement behavior parameters will depend strongly on whether the sampling interval is a (sub)multiple of the animal’s activity cycles: locations taken every 6 hours may provide highly biased estimates of home range size, speed, and diffusion if the animal of interest returns to its den every 6 hours to rest. This would be particularly visible in the unlucky case that all locations are taken when the animal is resting at its den and ignore the movement between each rest. For deer, the oscillations in speed and diffusion estimates seem to occur on a 12-hour cycle with peak mean diffusion at 12-hour and 36-hour sampling intervals and lower mean diffusion at sampling intervals of 24 and 48 hours. These findings may correlate with deer’s tendency to be less active during certain periods of the day (Roden-Reynolds et al., 2022). Still, these results further demonstrate the robustness of diffusion rate over speed in terms of both effective sample size and estimate stability, supporting the use of diffusion rate as a reliable metric for analyzing movement in coarse or irregular telemetry datasets. An interesting perspective would be to combine location data with activity data and read variations of mean speed and diffusion rates against diel activity cycles (Hertel et al., 2017).

### 4.3 Relationship between speed and diffusion rate in simulated trajectories

The results from the simulated telemetries supported the empirical findings while also demonstrating why estimating speed is not always appropriate. Forcing a model to estimate speed when telemetry data do not contain (sufficient) information on an animal’s speed and movement trajectory results in estimates that are most often biasedly low. This occurs because thinning telemetry data removes crucial fine-scale resolution on the tortuosity of the movement. While this is also true for diffusion, diffusion estimates are more reliant on the spatial variance of the locations (i.e., the area they cover) than the movement path.

While the results from the simulations strongly agree with the empirical results, they also demonstrate an important difference between the timescales of speed and diffusion. Since animals will change direction more often than they change their diffusion rate, speed changes at a faster rate than diffusion rate. Consequently, very fine sampling intervals will produce a large effective sample size for speed, but they will have a lesser impact on effective sample size for diffusion. This is because fine-scale sampling allows continuous-time models such as those fit by ctmm to obtain many independent estimates of speed (Noonan et al., 2019), but the number of independent diffusion values will be lower and often unaffected by sampling interval since it changes over longer timescales. Thus, if one is interested in estimating fine-scale, instantaneous estimates of speed (rather than simply mean speed), we echo the suggestion of Noonan et al. (2019) of choosing a sampling interval that is substantially (≥ 3 times) smaller than the directional persistence of the species of interest.

### 4.4 Implications and future research

Our results emphasize the complementary role of speed and diffusion rate in understanding animal movement and highlight the robustness and stability of diffusion rate when data quality is compromised due to coarse sampling or large measurement errors. In the absence of reasonable speed estimates, diffusion rate can serve as an effective alternative that allows inferences about movement patterns, space use, and energetics. By integrating both metrics, researchers can gain a more comprehensive understanding of movement behavior by using speed to assess fine-scale activity and energetics and diffusion to explore large-scale spatial patterns and ecological processes. This complementary approach allows researchers to infer behavioral and ecological dynamics even in challenging datasets and proposes diffusion rate as an essential tool for movement ecology alongside traditional metrics like mean speed and distance traveled.

However, the nuances in the relationship between diffusion and speed highlight the need for continued research into how diffusion relates to movement energetics and activity budgets. Understanding these dynamics could enhance the ecological insights derived from metrics of diffusion, especially across species and movement contexts, for example, complementing location and accelerometry data (see also dead-reckoning approaches, e.g., Magowan et al., 2022). By leveraging both metrics in future studies, researchers can bridge the gap between fine-scale behaviors and larger-scale ecological processes while achieving a more holistic view of animal movement and its implications for individuals and the surrounding community and landscape.

## Supporting information

Supplement A - Derivations

Supplement B - Estimating speed and diffusion from empirical data - method description

Supplement C - Interpreting the relationship between speed and diffusion

## DATA AVAILABILITY STATEMENT

The data and code used in this study are publicly available on the Open Science Framework (OSF) repository at the following link: https://osf.io/4hgev?view_only=c6ac7024b1454cffb662db8c5d79c3aa. This includes all datasets and analytical scripts necessary to reproduce the results presented in this manuscript.

## SUPPLEMENTARY FILES

Supplement A - Derivations

Supplement B - Estimating speed and diffusion from empirical data - method description

Supplement C - Interpreting the relationship between speed and diffusion

## AUTHOR CONTRIBUTIONS

Vickie DeNicola and Christen Fleming conceived the study, with the support of Francesca Cagnacci; Christen Fleming, Vickie DeNicola, and Stefano Mezzini designed the analytical study; Vickie DeNicola and Francesca Cagnacci designed the empirical study supporting the analytical study; Vickie DeNicola and Francesca Cagnacci led funding acquisition, project administration, and resource acquisition; Vickie DeNicola led the fieldwork and data collection; Vickie DeNicola and Stefano Mezzini curated the data; Stefano Mezzini and Vickie DeNicola led the development of the study methodology with support from Christen Fleming and Francesca Cagnacci; Stefano Mezzini led the formal analysis with support from Vickie DeNicola and Christen Fleming; Stefano Mezzini led the software development, design and interpretation of the models, and visualization of the data with support from Vickie DeNicola; Christen Fleming lead the mathematical theory and software implementation; Vickie DeNicola and Stefano Mezzini drafted the manuscript with support from Christen Fleming and Francesca Cagnacci. All authors assisted in revising the manuscript and gave final approval for publication. Christen Fleming and Francesca Cagnacci provided supervision.

