## Supplement A - Derivations for "Are your data too coarse for speed estimation? Diffusion rates as an alternative measure of animal movement"

### 1 Derivations

To compare the diffusion rates of different, selected stochastic process models, we first consider time-lag dependence of diffusion via the semi-variance function (SVF):

$$\gamma(\tau) = \frac{1}{4}\mathrm{E}\big[(x(t{+}\tau){-}x(t))^2 + (y(t{+}\tau){-}y(t))^2\big] \,, \tag{1.1}$$

which is a quarter of the time-averaged mean square displacement (MSD), defined so that $\gamma(\tau)$ limits to the average variance:

$$\lim_{\tau\to\infty} \gamma(\tau) = \sigma_0 = \frac{1}{2}\left(\mathrm{VAR}[x(t)] + \mathrm{VAR}[y(t)]\right) , \tag{1.2}$$

for a stationary movement process, and that

$$\gamma(\tau) = \sigma(0) - \sigma(\tau) \,, \tag{1.3}$$

in terms of the autocorrelation function (ACF), $\sigma(\tau)$.

The autocorrelation and semi-variance functions for our continuous-time stochastic process models are given in Fleming et al. (2015, 2017). From the SVF perspective, the diffusion rate can be defined in two different ways:

$$D(\tau) \equiv \frac{d\gamma}{d\tau}(\tau) = -\frac{d\sigma}{d\tau}(\tau) \,, \qquad \text{or} \qquad D(\tau) \equiv \frac{\gamma(\tau)}{\tau} \,, \tag{1.4}$$

which both measure how the MSD increases with increasing time lag. To compare diffusion rates across different stochastic process models, we will compare the maximum diffusion rates:

$$D_{\mathrm{max}} = D(\tau_{\mathrm{max}}) \qquad \text{where} \qquad \tau_{\mathrm{max}} = \arg\max_{\tau} D(\tau) \,. \tag{1.5}$$

As we will show, this definition of the maximum diffusion rate produces the commonly accepted parameters of comparison between Brownian motion (BM), Ornstein-Uhlenbeck(OU), and integrated Ornstein-Uhlenbeck (IOU) processes. Furthermore, we will use the first definition in (1.4), which relies on differentiation, because our parameters of interest can then be solved in closed form, from either $\gamma''(\tau_{\mathrm{max}}) = 0$ or $\sigma''(\tau_{\mathrm{max}}) = 0$. However, while all of the SVFs we consider are monotonically increasing functions, the second diffusion-rate definition in (1.4) would be advantageous if that were not the case, as the maximum slope might not be particularly meaningful then.

#### 1.1 Brownian motion

The Brownian motion SVF is simply linear

$$\gamma(\tau) = D\,\tau \tag{1.6}$$

with the maximum diffusion rate given by the slope, $D$, which is the only diffusion rate for this process.

#### 1.2 Ornstein-Uhlenbeck motion

The Ornstein-Uhlenbeck ACF is given by:

$$\sigma(\tau) = \sigma_0\, e^{-\frac{\tau}{\tau_p}} \tag{1.7}$$

in terms of the position autocorrelation timescale, $\tau_p$. The maximum diffusion rate is then given by the instantaneous diffusion rate:

$$D_0 = \frac{\sigma_0}{\tau_p}\,. \tag{1.8}$$

#### 1.3 Integrated Ornstein-Uhlenbeck motion

The integrated Ornstein-Uhlenbeck SVF is given by:

$$\gamma(\tau) = D_\infty \left(\tau - \tau_v \left(1 - e^{-\frac{\tau}{\tau_v}}\right)\right), \tag{1.9}$$

in terms of the velocity autocorrelation timescale, $\tau_v$. The maximum diffusion rate given by the asymptotic diffusion rate, $D_\infty$.

#### 1.4 OUF motion

The OUF ACF is given by (Fleming et al., 2014):

$$\gamma(\tau) = \sigma_0 \frac{\tau_p\, e^{-\frac{\tau}{\tau_p}} - \tau_v\, e^{-\frac{\tau}{\tau_v}}}{\tau_p - \tau_v}. \tag{1.10}$$

By differentiating the ACF twice, we find the maximum diffusion rate to occur at time lag

$$\tau_{\mathrm{max}} = \frac{\log\theta}{\theta-1}\,\tau_p\,, \qquad \text{where} \qquad \theta \equiv \frac{\tau_p}{\tau_v}\,, \tag{1.11}$$

and by plugging this back into the derivative of the SVF, we find the maximum diffusion rate to be:

$$D_{\mathrm{max}} = \frac{1}{\sqrt[\theta-1]{\theta}}\,\frac{\sigma_0}{\tau_p}\,. \tag{1.12}$$

#### 1.5 OUO motion

The oscillatory OUF ACF—termed OUO in `ctmm`—is given by (Fleming et al., 2017):

(1.13)

$$\gamma(\tau) = \sigma_0 \left( \cos(\omega\tau) + \frac{\sin(\omega\tau)}{\omega\tau_p} \right) e^{-\frac{\tau}{\tau_p}} \, ,$$

in terms of the oscillation frequency, $\omega$. Again, by differentiating the ACF twice, we find the maximum diffusion rate to occur at time lag

$$\tau_{\text{max}} = \frac{\tan^{-1} \epsilon}{\epsilon} \, \tau_p \, , \qquad \text{where} \qquad \epsilon \equiv \omega\tau_p \, , \tag{1.14}$$

and by plugging this back into the derivative of the SVF, we find the maximum diffusion rate to be:

$$D_{\text{max}} = \sqrt{1+\epsilon^2} \, e^{-\frac{\tan^{-1}\epsilon}{\epsilon}} \, \frac{\sigma_0}{\tau_p} \, . \tag{1.15}$$

#### 1.6 OUf motion

The OUf ACF, which is OUF where $\tau_p = \tau_v$ or OUO where $\omega = 0$, is given by:

$$\gamma(\tau) = \sigma_0 \left( 1 + \frac{\tau}{\tau_p} \right) e^{-\frac{\tau}{\tau_p}} \, . \tag{1.16}$$

The maximum diffusion rate is easily obtained from the $\omega \to 0$ limit of the OUO result:

$$D_{\text{max}} = \frac{1}{e} \, \frac{\sigma_0}{\tau_p} \, . \tag{1.17}$$
