## Supplement B - Estimating speed and diffusion from empirical data - method description for "Are your data too coarse for speed estimation? Diffusion rates as an alternative measure of animal movement"

Estimating speed and diffusion from empirical data – reproducing the analyses

```
library('dplyr')   # for data wrangling
library('tidyr')   # for data wrangling
library('purrr')   # for functional programming
library('furrr')   # for parallelized functional programming
library('ggplot2') # for fancy plots
library('mecor')   # for models with measurement error correction
library('ctmm')    # for movement models
library('mgcv')    # for generalized additive models
library('gratia')  # for ggplot-based diagnostics for GAMs
library('khroma')  # for colorblind-friendly color palettes
library('ggExtra') # for marginal density plots and histograms
library('cowplot') # for fancy multi-panel plots
```

### 1 Relationship between mean speed and diffusion rate (Fig.1)

```r
source('analysis/figures/default-ggplot-theme.R') # gets reset in new chunk
COL <- '#7FB0E3' # color for estimated relationship and CIs

d <- readRDS('../ny-deer-vasectomy/data/years-1-and-2-data-no-akde.rds') %>%
  filter(is.finite(speed_est)) %>% # loosing ~42% of the windows
  # drop animals with median sampling intervals > 4-hours
  mutate(dt_hours = map_dbl(tel, \(.t) {
    round(as.numeric(median(diff(.t$timestamp)), units = 'hours'))
  })) %>%
  filter(dt_hours == 1) %>%
  #' fix units ("square kilometers/day" is not recognized by `%#%`)
  mutate(diffusion_units = 'km^2/day') %>%
  select(animal, animal_year, tel, speed_est, diffusion_est, dof_speed,
         dof_diff, speed_units, diffusion_units, dt_hours) %>%
  rename(dof_diffusion = dof_diff)

nrow(d) # number of 7-day speed estimates

## [1] 5180

n_distinct(d$animal_year) # total deer with duplicates between years

## [1] 117

n_distinct(gsub(' .*', '', d$animal_year)) # total deer w/o duplicates

## [1] 108

ggplot(d) +
  # data points
  geom_point(aes(diffusion_est, speed_est), alpha = 0.1) +
  # estimated relationship
  geom_smooth(aes(diffusion_est, speed_est), color = COL, method = 'lm',
              formula = y ~ x) +
  scale_x_continuous('Estimated diffusion (km\U00B2/day, log scale)',
                     transform = 'log', labels = \(x) round(x, 2)) +
  scale_y_continuous('Estimated speed (km/day, log scale)',
                     transform = 'log', labels = \(x) round(x, 2)) +
  ggtitle('Exploratory plot with geom_smooth')
```

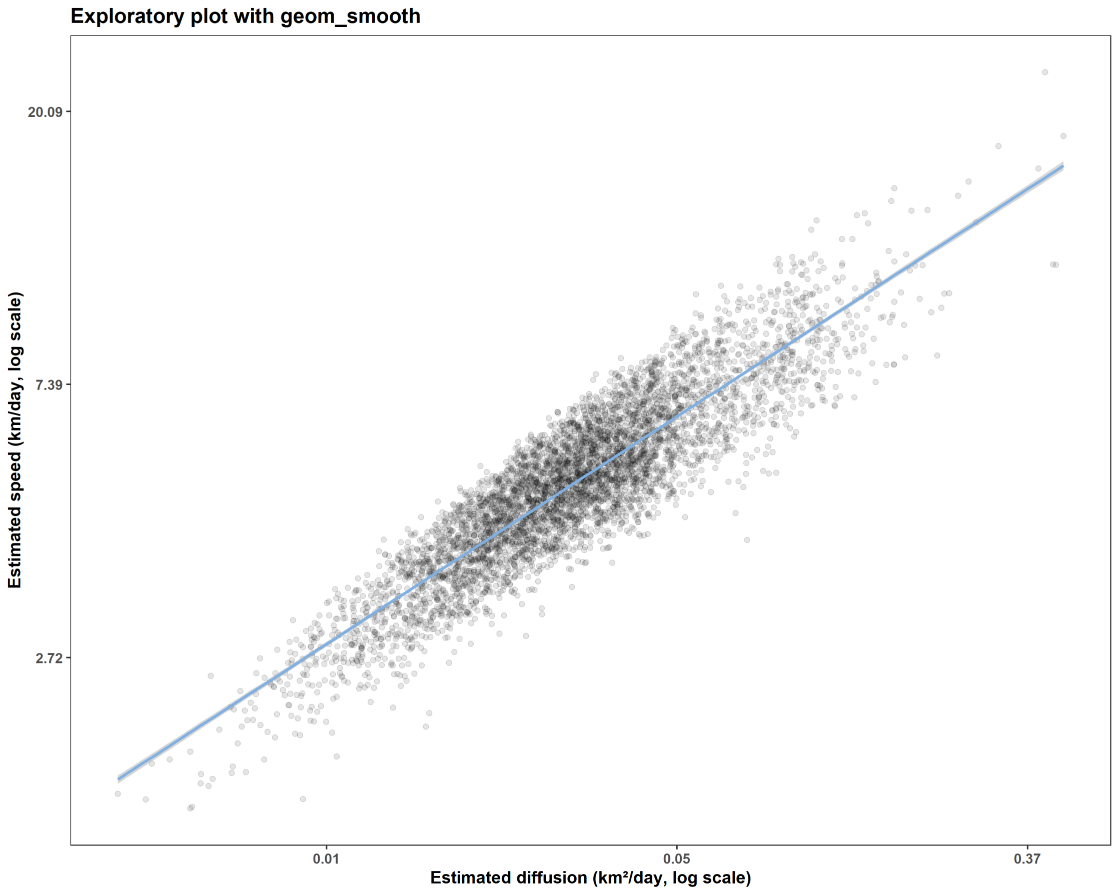


```r
# fit the model ----
# not using weights because they bias towards smaller tau values
m <- bam(speed_est ~
           log(diffusion_est) +
           s(log(diffusion_est), animal, bs = 'fs'),
         family = Gamma(link = 'log'), data = d, method = 'fREML',
         discrete = TRUE)

draw(m, parametric = TRUE)
```

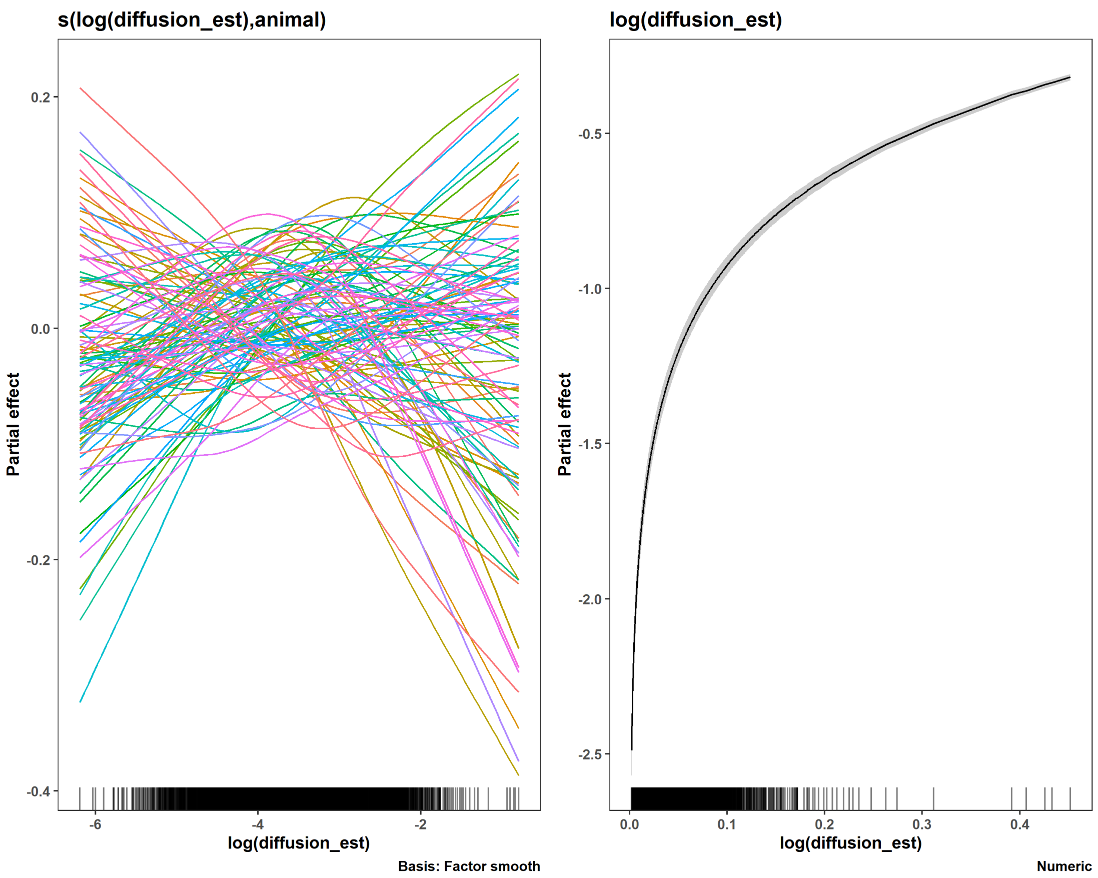


```
summary(m)

##
## Family: Gamma
## Link function: log
##
## Formula:
## speed_est ~ log(diffusion_est) + s(log(diffusion_est), animal,
##     bs = "fs")
##
## Parametric coefficients:
##                    Estimate Std. Error t value Pr(>|t|)
## (Intercept)        3.090445   0.024526  126.01   <2e-16 ***
## log(diffusion_est) 0.402003   0.006633   60.61   <2e-16 ***
## ---
## Signif. codes:  0 '***' 0.001 '**' 0.01 '*' 0.05 '.' 0.1 ' ' 1
##
## Approximate significance of smooth terms:
##                               edf Ref.df     F p-value
## s(log(diffusion_est),animal) 183.8   1011 1.031  <2e-16 ***
## ---
## Signif. codes:  0 '***' 0.001 '**' 0.01 '*' 0.05 '.' 0.1 ' ' 1
##
```

```
## R-sq.(adj) =  0.828   Deviance explained = 84.6%
## fREML = -3147.5  Scale est. = 0.016093  n = 5180

appraise(m, point_alpha = 0.1)
```

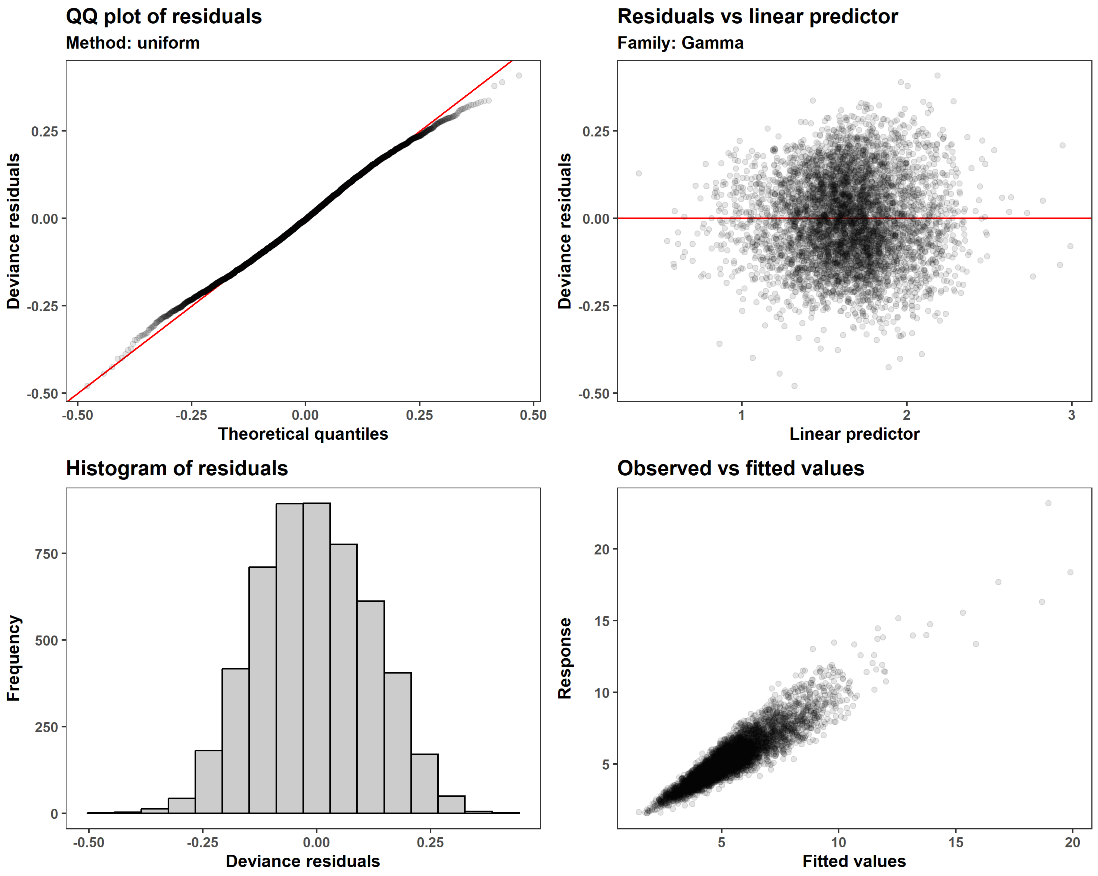


```
# est and CIs for slope of log(diffusion)
b1 <- round(coef(m)['log(diffusion_est)'], 4); b1

## log(diffusion_est)
##              0.402

round(b1 + summary(m)$se['log(diffusion_est)'] * qnorm(c(0.005, 0.995)), 4)

## [1] 0.3849 0.4191

# plot the model and data ----
preds <-
  tibble(diffusion_est = seq(1.5e-3, 0.6, length.out = 400),
         dof_speed = mean(d$dof_speed),
         animal = 'new animal') %>%
  bind_cols(
    .,
    predict(object = m, newdata = ., type = 'link', se.fit = TRUE,
            terms = c('log(diffusion_est)', '(Intercept)'),
```

```
            discrete = FALSE) %>%
      as.data.frame() %>%
      transmute(lwr_95 = exp(fit + qnorm(0.005) * se.fit),
                mu_hat = exp(fit),
                upr_95 = exp(fit + qnorm(0.995) * se.fit)))

## Warning in predict.gam(object, newdata = newdata, type = type, se.fit = se
.fit,
## : factor levels new animal not in original fit

preds

## # A tibble: 400 × 6
##    diffusion_est dof_speed animal     lwr_95 mu_hat upr_95
##  *         <dbl>     <dbl> <chr>       <dbl>  <dbl>  <dbl>
##  1        0.0015      25.9 new animal   1.53   1.61   1.69
##  2        0.003       25.9 new animal   2.05   2.13   2.21
##  3        0.0045      25.9 new animal   2.42   2.50   2.59
##  4        0.006       25.9 new animal   2.73   2.81   2.89
##  5        0.0075      25.9 new animal   3.00   3.08   3.16
##  6        0.009       25.9 new animal   3.23   3.31   3.39
##  7        0.0105      25.9 new animal   3.45   3.52   3.60
##  8        0.012       25.9 new animal   3.64   3.72   3.79
##  9        0.0135      25.9 new animal   3.83   3.90   3.97
## 10        0.015       25.9 new animal   4.00   4.06   4.13
## # ℹ 390 more rows

# fig. 1
ggplot() +
  # data points
  geom_point(aes(diffusion_est, speed_est), d, alpha = 0.1) +
  # estimated relationship
  geom_ribbon(aes(diffusion_est, ymin = lwr_95, ymax = upr_95), preds,
              alpha = 0.3, fill = COL) +
  geom_line(aes(diffusion_est, mu_hat), preds, color = COL, lwd = 1) +
  scale_x_continuous('Estimated diffusion (km\U00B2/day, log scale)',
                     transform = 'log', labels = \(x) round(x, 2)) +
  scale_y_continuous('Estimated speed (km/day, log scale)',
                     transform = 'log', labels = \(x) round(x, 2))
```

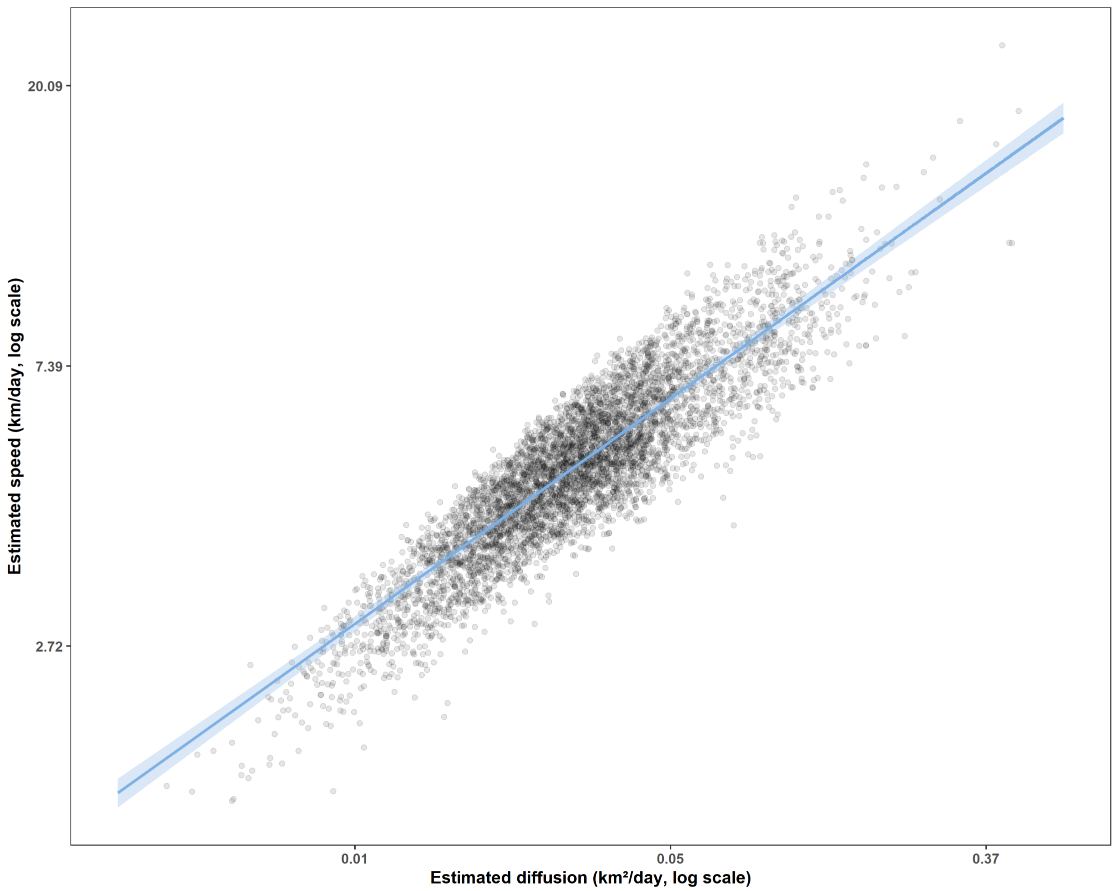


```r
# for graphical abstract
tibble(diffusion_est = seq(0, 0.6, length.out = 400),
       dof_speed = mean(d$dof_speed),
       animal = 'new animal') %>%
  bind_cols(
    .,
    predict(object = m, newdata = ., type = 'link', se.fit = TRUE,
            terms = c('log(diffusion_est)', '(Intercept)'),
            discrete = FALSE) %>%
      as.data.frame() %>%
      transmute(lwr_95 = exp(fit + qnorm(0.005) * se.fit),
                mu_hat = exp(fit),
                upr_95 = exp(fit + qnorm(0.995) * se.fit))) %>%
  ggplot() +
  # estimated relationship
  geom_ribbon(aes(diffusion_est, ymin = lwr_95, ymax = upr_95),
              alpha = 0.3, fill = COL) +
  geom_line(aes(diffusion_est, mu_hat), color = COL, lwd = 1) +
  scale_x_continuous('Estimated diffusion (km\U00B2/day)') +
  scale_y_continuous('Estimated speed (km/day)')
```

```
## Warning in predict.gam(object, newdata = newdata, type = type, se.fit = se
.fit,
## : factor levels new animal not in original fit
```

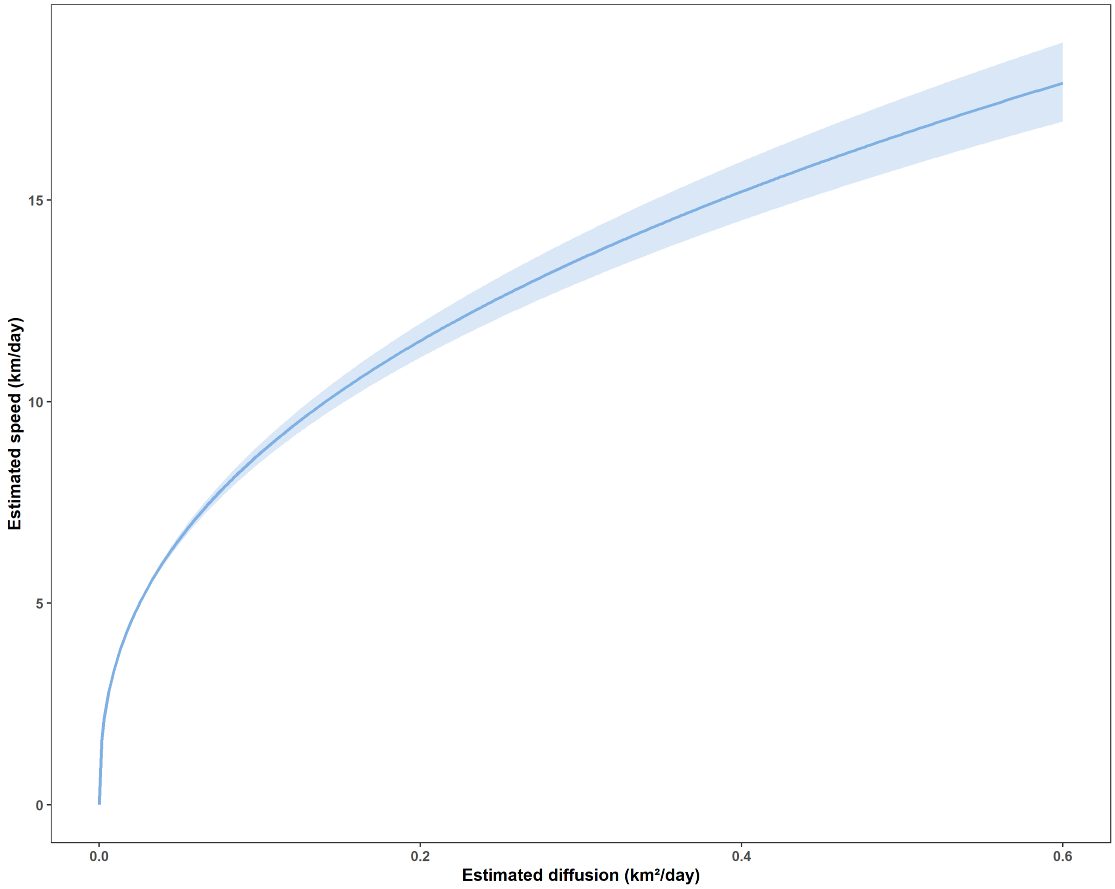


```
# estimated speed decreases with speed effective sample size (ess),
# but we don't have an estimate of the ess when speed is NA
ggplot(d, aes(log(dof_speed), residuals(m))) +
  geom_point(alpha = 0.1) +
  geom_smooth(color = 'darkorange', method = 'gam',
              formula = y ~ s(x, bs = 'ts', k = 4), n = 400) +
  labs(x = expression(bold(log(Estimated~speed~n[eff]))), y = 'Residuals')
```

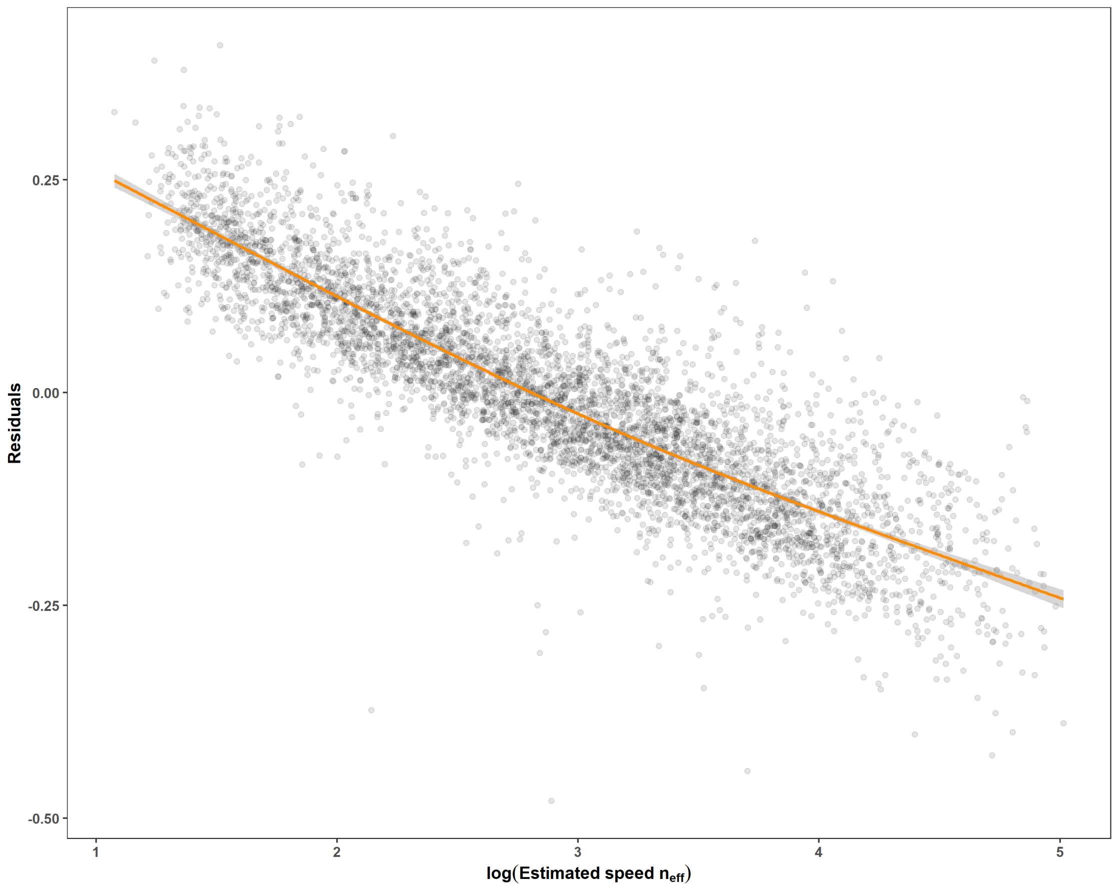


```
# some correlation between the dof for speed and for diffusion
ggplot(d, aes(log(dof_speed), log(dof_diffusion))) +
  geom_point(alpha = 0.1) +
  labs(x = expression(bold(log(Estimated~diffusion~n[eff]))),
       y = expression(bold(log(Estimated~speed~n[eff]))))
```

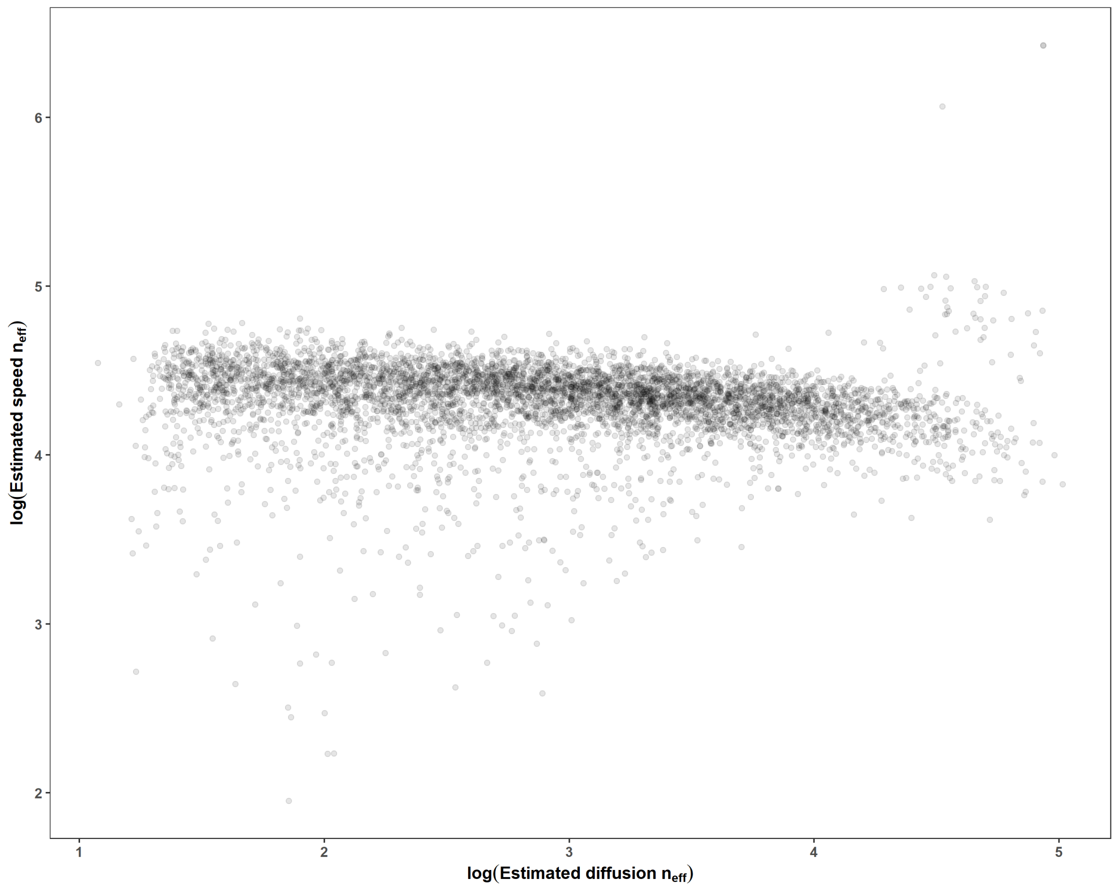


```r
# low DOF are more likely to under-estimate diffusion
# this may be due to the correlation between speed and diffusion
ggplot(d, aes(log(dof_diffusion), residuals(m))) +
  geom_smooth(color = 'darkorange', method = 'gam',
              formula = y ~ s(x, bs = 'ts', k = 4), n = 400) +
  geom_point(alpha = 0.1) +
  labs(x = expression(bold(log(Estimated~diffusion~n[eff]))),
       y = 'Residuals')
```

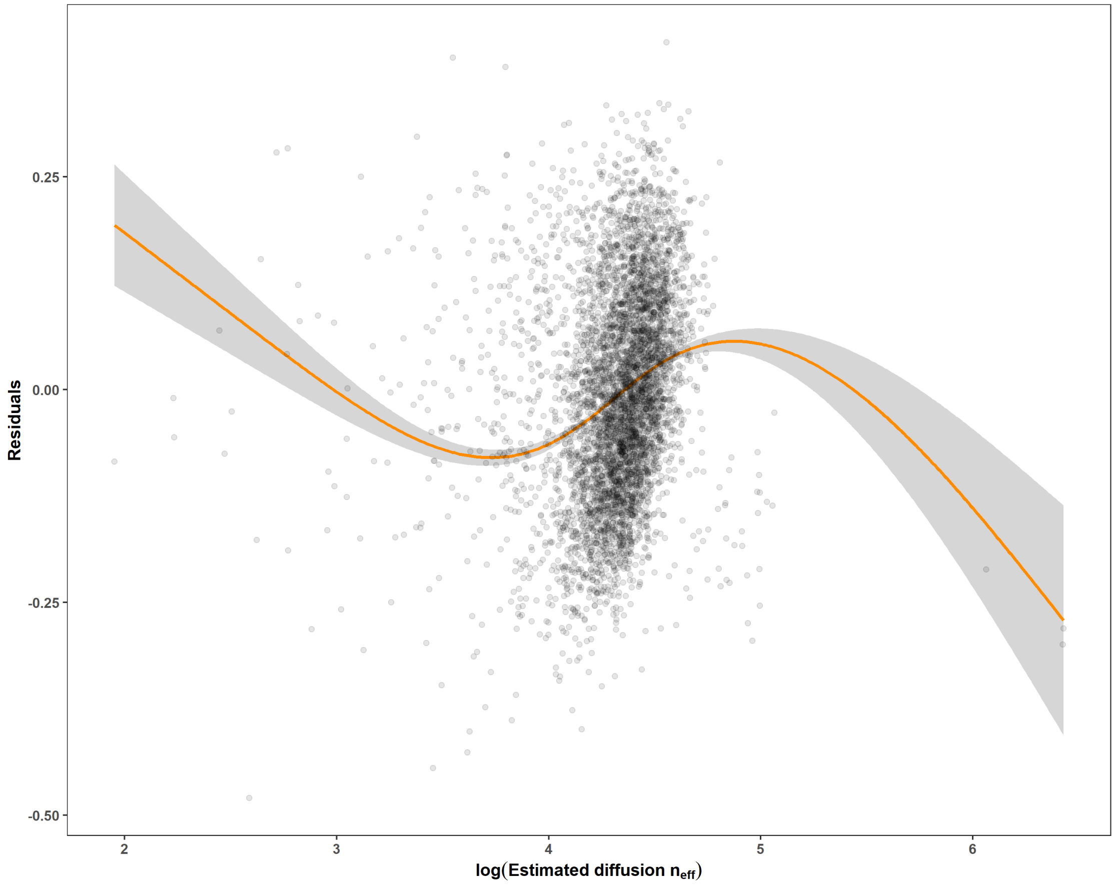


```r
#' same plot but for `4 < log(dof_diffusion) < 5`
d %>%
  mutate(e = residuals(m)) %>%
  filter(log(dof_diffusion) > 4, log(dof_diffusion) < 5) %>%
  ggplot(aes(log(dof_diffusion), e)) +
  geom_smooth(color = 'darkorange', method = 'gam',
              formula = y ~ s(x, bs = 'ts', k = 4), n = 400) +
  geom_point(alpha = 0.1) +
  labs(x = expression(bold(log(Estimated~diffusion~n[eff]))),
       y = 'Residuals')
```

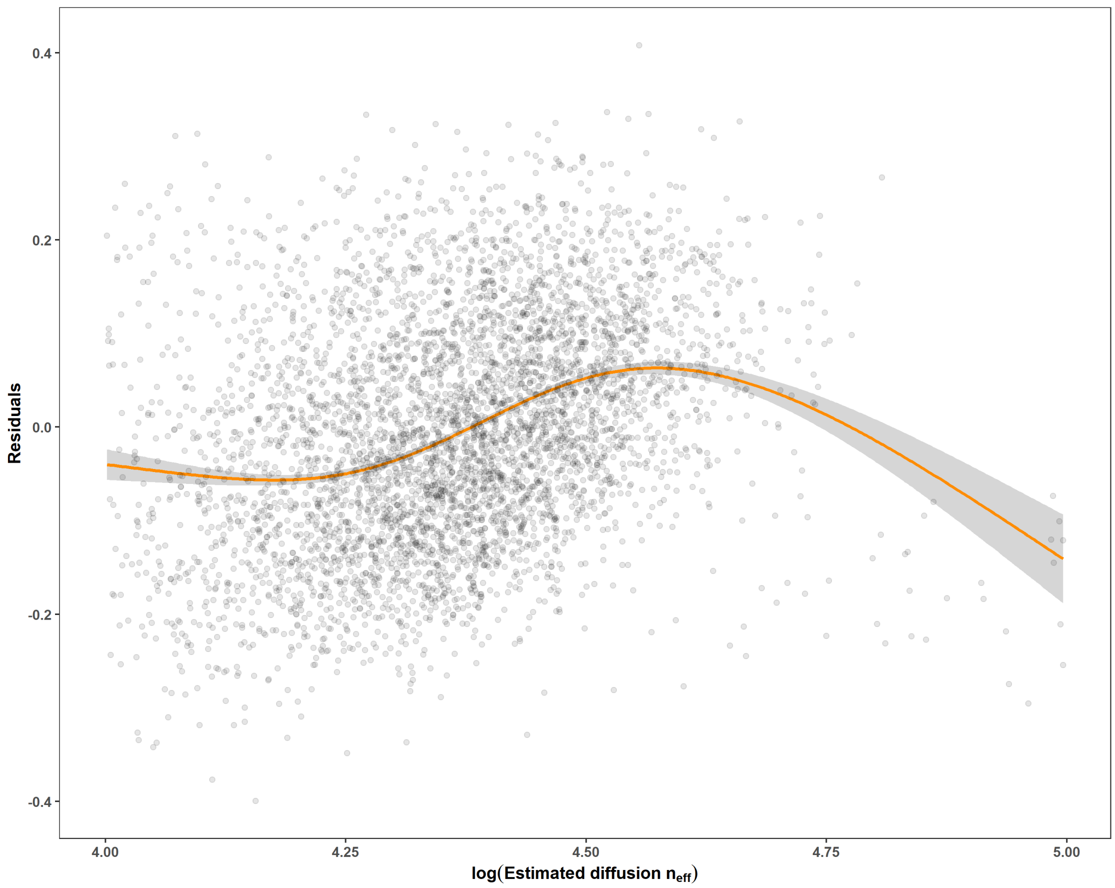
0.4
0.2
0.0
-0.2
-0.4
Residuals
4.00
4.25
4.50
4.75
5.00
log(Estimated diffusion $n_{eff}$)

```r
source('analysis/figures/default-ggplot-theme.R') # gets reset in new chunk
tels <-
  bind_rows(
    data.table::fread('data/deer-data/Year_1_processed_data.csv'),
    data.table::fread('data/deer-data/Year_2_processed_data.csv')) %>%
  # fix column names to avoid needing backticks
  rename_with(\(.x) gsub('-', '_', .x)) %>%
  rename_with(\(.x) gsub(':', '_', .x)) %>%
  rename_with(\(.x) gsub('\\.', '_', .x)) %>%
  # only keeep necessary columns
  select(c(event_id, timestamp, location_long, location_lat, gps_dop,
           gps_fix_type_raw, gps_satellite_count,
           individual_taxon_canonical_name, individual_local_identifier,
           Date, Time, Year, Month, Day, study_area, animal_sex,
           study_year, animal_age)) %>%
  # add a duplicate animal name for nesting telemetries later
  mutate(animal = individual_local_identifier) %>%
  # get intervals between locations
  group_by(animal, study_year) %>%
  mutate(dt_hours = 'hours' %#% c(NA_real_, diff(timestamp))) %>%
  ungroup()

# find number of individuals
tels %>%
  group_by(animal) %>%
  slice(1) %>%
  pull(animal_sex) %>%
  table(dnn = 'Total deer by sex')

## Total deer by sex
##  f  m
## 76 38

# check range of time between locations
range(round(tels$dt_hours, 4), na.rm = TRUE)

## [1]  0.0004 25.2249

ggplot(tels, aes(dt_hours)) +
  geom_histogram(binwidth = 0.5, center = 0) +
  scale_y_continuous(trans = 'sqrt')

## Warning: Removed 124 rows containing non-finite outside the scale range
## (`stat_bin()`).
```

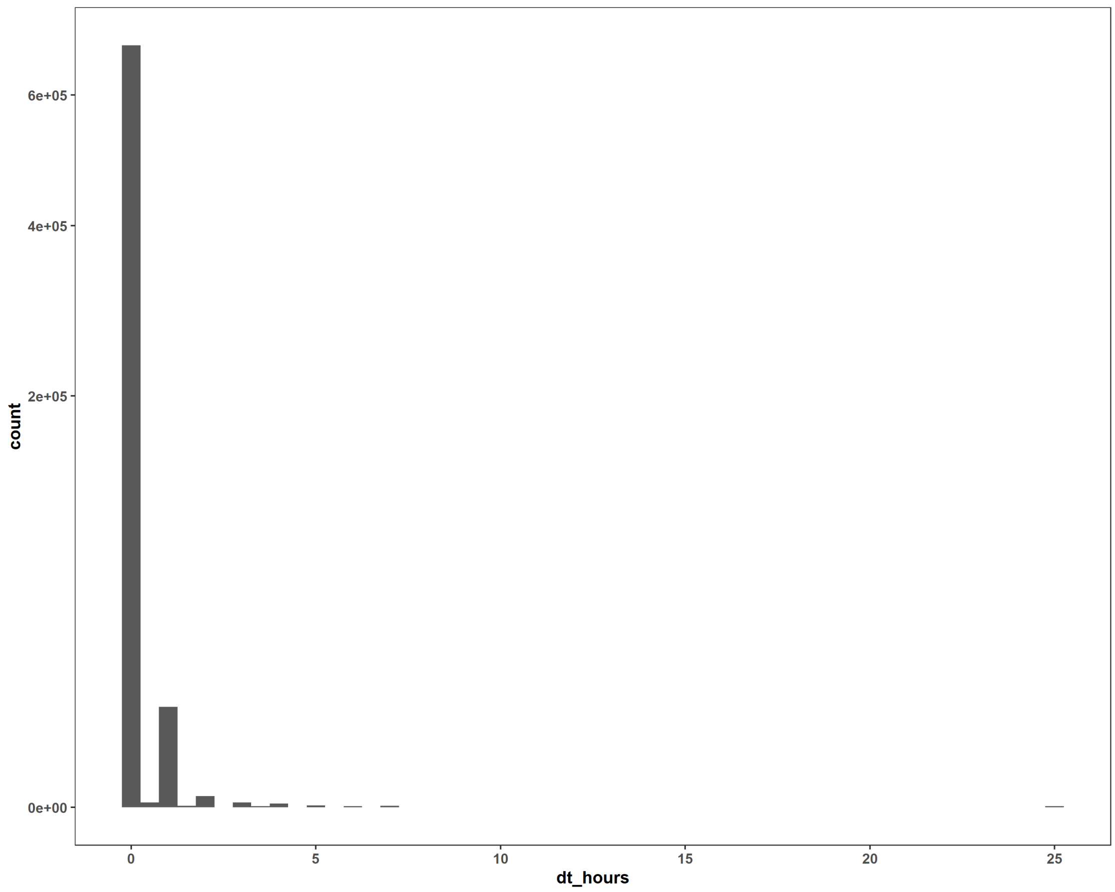


```r
ggplot(tels, aes(dt_hours)) +
  coord_cartesian(ylim = c(0, 50)) +
  geom_histogram()
```

```
## `stat_bin()` using `bins = 30`. Pick better value with `binwidth`.
```

```
## Warning: Removed 124 rows containing non-finite outside the scale range
## (`stat_bin()`).
```

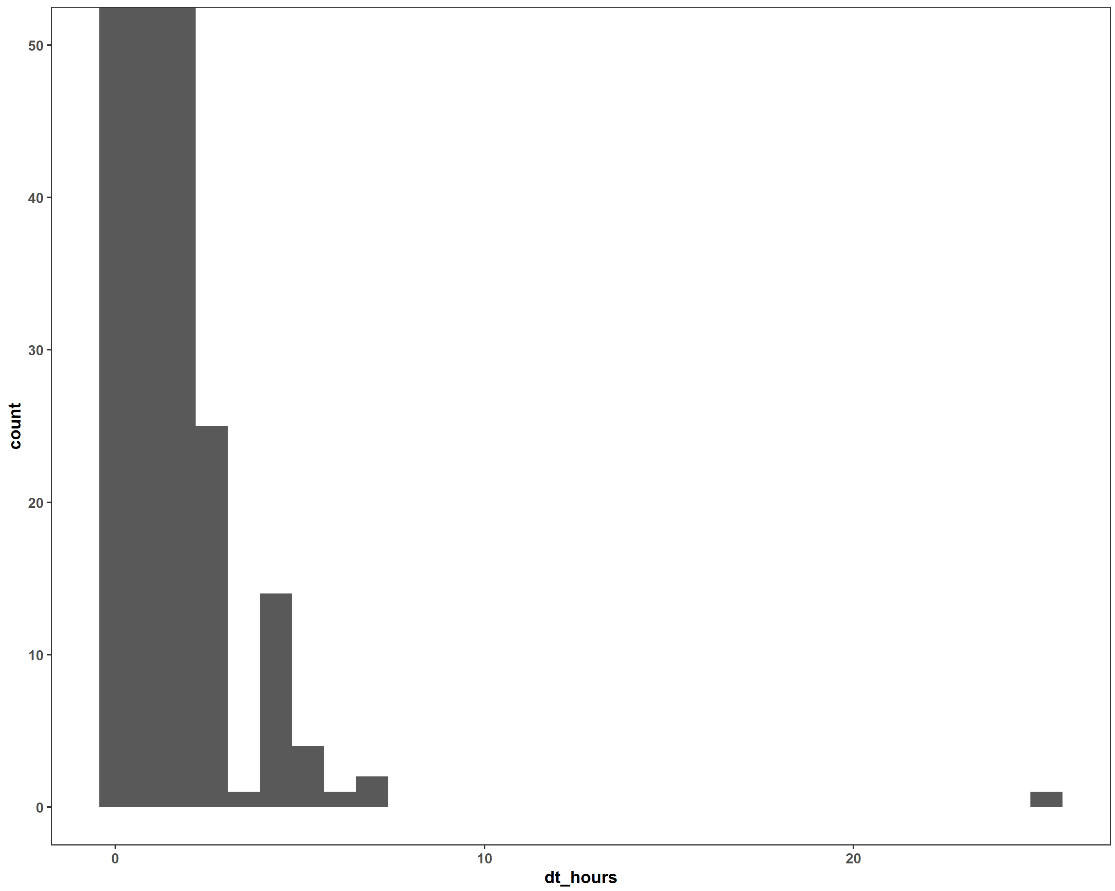


```
thinned <- tibble(
  dt_hours = seq(1, 48, by = 1),
  data = map(dt_hours, function(.dt) {
    tels %>%
      group_by(animal, study_year) %>%
      filter(
        round('hours'%#%as.numeric(timestamp-min(timestamp)))%%.dt == 0) %>%
      ungroup()
  }),
  n_total = map_int(data, nrow))
thinned
```

```
## # A tibble: 48 × 3
##    dt_hours data                      n_total
##       <dbl> <list>                      <int>
##  1        1 <tibble [698,813 × 20]>    698813
##  2        2 <tibble [351,681 × 20]>    351681
##  3        3 <tibble [232,834 × 20]>    232834
##  4        4 <tibble [178,554 × 20]>    178554
##  5        5 <tibble [139,761 × 20]>    139761
##  6        6 <tibble [117,232 × 20]>    117232
##  7        7 <tibble [99,859 × 20]>      99859
##  8        8 <tibble [89,331 × 20]>      89331
```

```
## 9         9 <tibble [77,662 × 20]>    77662
## 10       10 <tibble [70,341 × 20]>    70341
## # ℹ 38 more rows
```

```
ggplot(thinned, aes(dt_hours, n_total)) +
  geom_line(aes(y = nrow(tels) / dt_hours), col = 'grey') +
  geom_point(alpha = 0.5) +
  geom_hline(yintercept = nrow(tels), lty = 'dashed') +
  xlab('Sampling interval') +
  scale_y_log10('Number of locations', limits = c(1e4, NA))
```

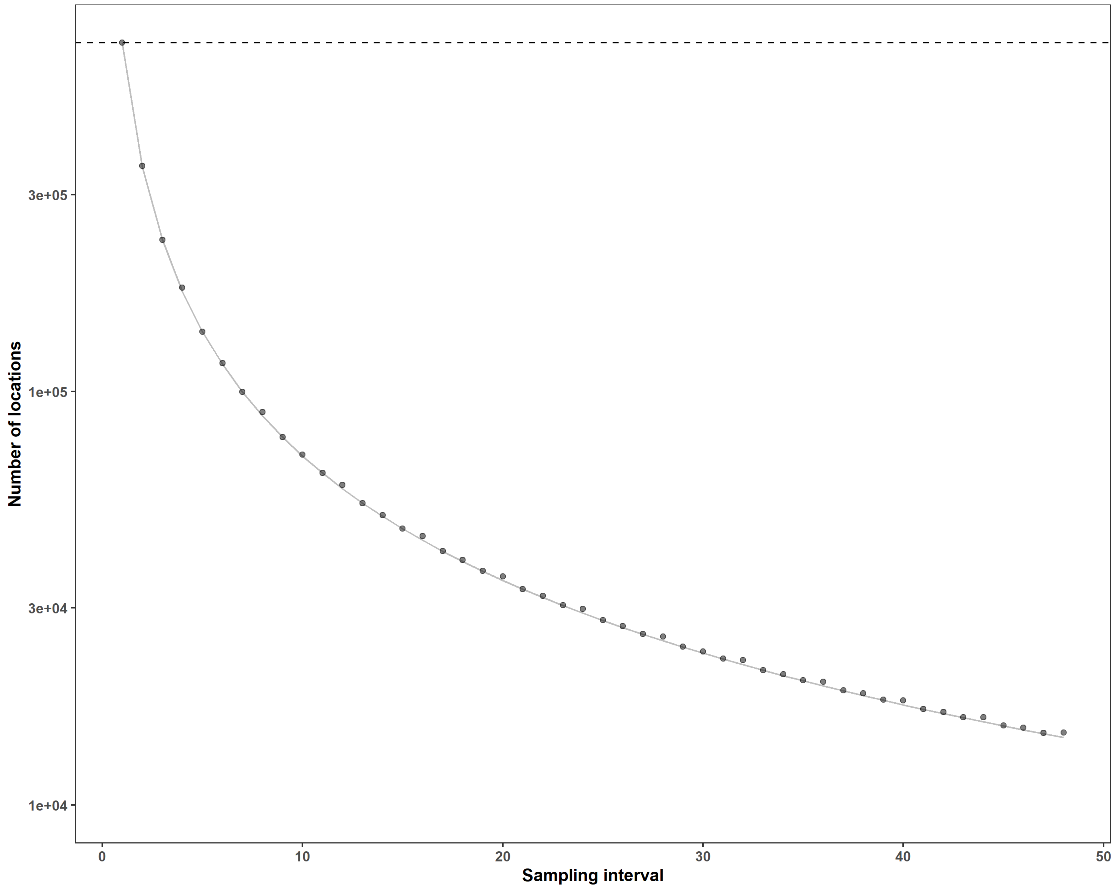

#### 3 Fitting movement models to the thinned telemetries

To avoid unnecessary computations, we do not run this chunk while knitting the pdf (using eval = FALSE). The thinned-models-dt-*-hours.rds files are on GitHub and can be merged using the map_dfr call at the end of the code chunk.

```
NCORES <- min(12, availableCores(logical = FALSE) - 2)
plan(multisession, workers = NCORES)

START <- Sys.time()
START

d0 <- readRDS('data/deer-data/thinned-telemetry-data.rds')

#' if you get an error about an NA value in the `ctmm.select()` call,
#' restart R and run the code again
map(nrow(d0):1, function(i) {
  i <<- i

  d0 %>%
    slice(i) %>%
    mutate(data = map(data, \(.d) select(.d, ! dt_hours))) %>%
    unnest(data) %>%
    nest(tel = ! c(animal, dt_hours, study_area, study_year, animal_sex,
                   animal_age, n_total)) %>%
    mutate(tel = map(tel, \(.tel) suppressMessages(as.telemetry(.tel))),
           vg = map(tel, \(.tel) ctmm.guess(.tel, interactive = FALSE)),
           mm = future_map2(tel, vg, \(.tel, .vg) {
             ctmm.select(data = .tel, CTMM = .vg)
           }, .options = furrr_options(seed = NULL), .progress = TRUE)) %>%
    mutate(speed = map(mm, \(.mm) {
      x <- suppressWarnings(speed(.mm, units = FALSE))

      if(x$DOF == 0) {
        tibble(speed_dof = 0,
               speed_lwr_95 = 0,
               speed_est = Inf,
               speed_upr_95 = Inf) %>%
          return()
      } else {
        bind_cols(speed_dof = x$DOF,
                  as.data.frame('km/day' %#% x$CI) %>%
                    rename(speed_lwr_95 = low,
                           speed_est = est,
                           speed_upr_95 = high)) %>%
          return()
      }
    }),
    diff_dof = map_dbl(mm, \(.mm) summary(.mm,units=FALSE)$DOF['diffusion']),
```

```r
    diff_lwr = 'km^2/day' %#% map_dbl(mm, \(.m) ctmm:::diffusion(.m)[1]),
    diff_est = 'km^2/day' %#% map_dbl(mm, \(.m) ctmm:::diffusion(.m)[2]),
    diff_upr = 'km^2/day' %#% map_dbl(mm, \(.m) ctmm:::diffusion(.m)[3])) %>%
    unnest(speed) %>%
    saveRDS(paste0('models/deer-models/thinned-models-dt-', d0$dt_hours[i],
                   '-h.rds'))
  END <- Sys.time()
  return(paste(i, END))
})
beepr::beep()

# merge all files into one
map_dfr(1:48, \(.h) paste0('models/deer-models/thinned-models-dt-', .h,
                           '-h.rds') %>%
          readRDS()) %>%
  saveRDS('models/deer-models/thinned-movement-models.rds')
```

#### 4 Estimating speed and diffusion from the movement models (Fig.2)

```r
source('analysis/figures/default-ggplot-theme.R') # gets reset in new chunk
d <- readRDS('models/deer-models/thinned-movement-models.rds') %>%
  mutate(animal_year = paste(animal, study_year),
         duration = map_dbl(tel, \(.t) {
           diff(range(.t$timestamp))
         }),
         dt_hours_actual = map_dbl(tel, \(.t) {
           as.numeric(median(diff(.t$timestamp)), units = 'hours')
         })) %>%
  select(! n_total:mm)

# we start with 114 animals and 124 telemetries (10 tracked twice)
n_distinct(d$animal)

## [1] 114

n_distinct(d$animal_year)

## [1] 124

nrow(d) / 48 # should be 124 total telemetries

## [1] 124

DROP <- d %>%
  filter(dt_hours == 1) %>%
  group_by(animal_year) %>%
  summarise(difference = median(dt_hours_actual - dt_hours)) %>%
  filter(abs(difference) > 1)

hist(DROP$difference)
```

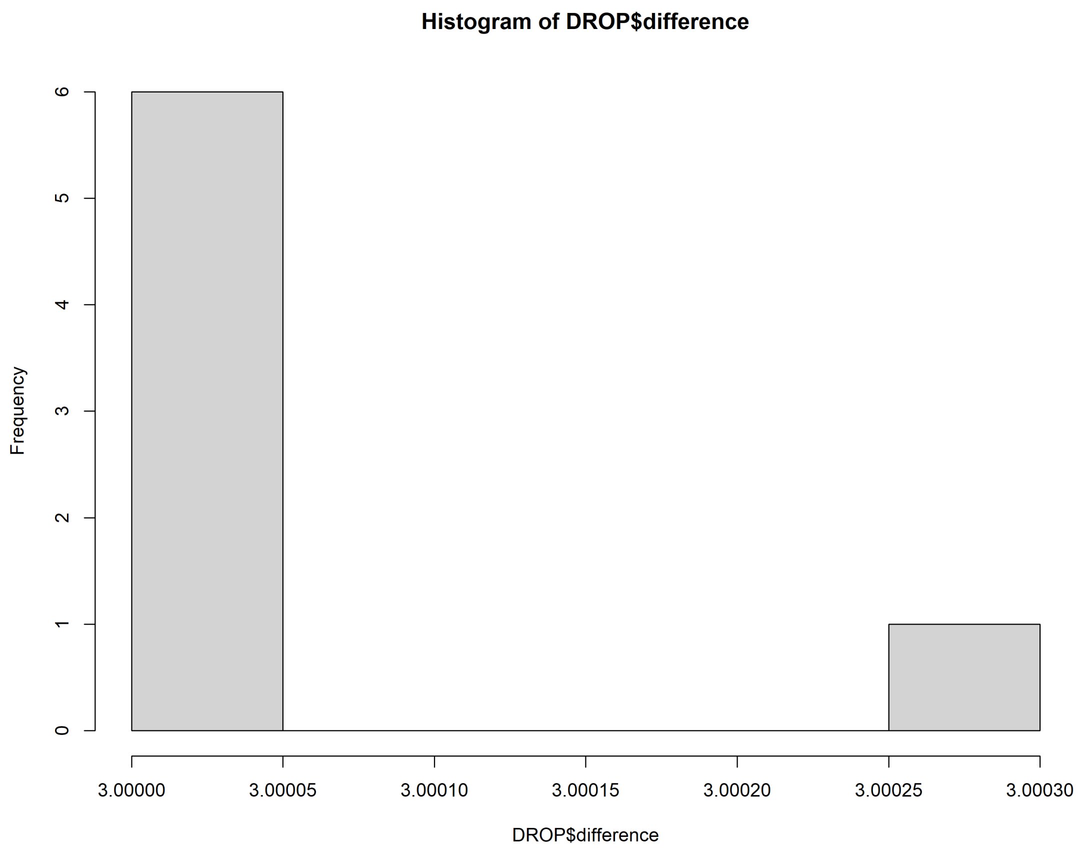


```r
#' drop animals with sampling interval that does not match `dt_hours`
d <- filter(d, ! animal_year %in% DROP$animal_year)

nrow(d) / 48

## [1] 117

n_distinct(d$animal_year) # lost 7 telemetries

## [1] 117

n_distinct(d$animal) # lost 6 animals

## [1] 108

d <- d %>%
  mutate(animal_year = factor(paste(animal, study_year))) %>%
  group_by(animal_year) %>%
  mutate(weight = sqrt(speed_dof) / mean(sqrt(speed_dof))) %>%
  ungroup()

# percenge of models with finite speed and diffusion
d %>%
  summarise(n = n(),
```

```
            diffusion = sum(is.finite(d$diff_est)),
            speed = sum(is.finite(d$speed_est)),
            both = sum(is.finite(d$diff_est) & is.finite(d$speed_est))) %>%
  mutate(p_diffusion = diffusion / n * 100,
         p_speed = speed / n * 100,
         p_both = both / n * 100)

## # A tibble: 1 × 7
##       n diffusion speed  both p_diffusion p_speed p_both
##   <int>     <int> <int> <int>       <dbl>   <dbl>  <dbl>
## 1  5616      4219   984   984        75.1    17.5   17.5

# check number of models per sampling interval ----
#' There are too few deer to subset to only deer contained in all sampling
#' intervals. Ideally we would have many deer with movement models for all
#' sampling intervals, but we have so few that we should model all deer.
nrow(d) / n_distinct(d$animal)

## [1] 52

n_models <- d %>%
  group_by(animal, study_year) %>%
  summarise(n = n())

## `summarise()` has grouped output by 'animal'. You can override using the
## `.groups` argument.

#' `DROP` dropped some animals entirely
nrow(n_models) # was 117 before dropping NAs and

## [1] 117

n_distinct(d$animal) # should be 106

## [1] 108

d %>%
  select(animal, animal_year, dt_hours, dt_hours_actual)

## # A tibble: 5,616 × 4
##    animal animal_year dt_hours dt_hours_actual
##    <chr>  <fct>          <dbl>           <dbl>
##  1 148    148 1              1            1.00
##  2 1074   1074 1             1            1
##  3 3      3 1                1            1
##  4 2      2 1                1            1.00
##  5 629    629 1              1            1
##  6 20a    20a 1              1            1
##  7 11     11 1               1            1
##  8 12     12 1               1            1
##  9 1      1 1                1            1
```

```
## 10 6a     6a 1                 1             1
## # ℹ 5,606 more rows
```

```r
ggplot(d, aes(dt_hours_actual, dt_hours)) +
  geom_point(alpha = 0.03)
```

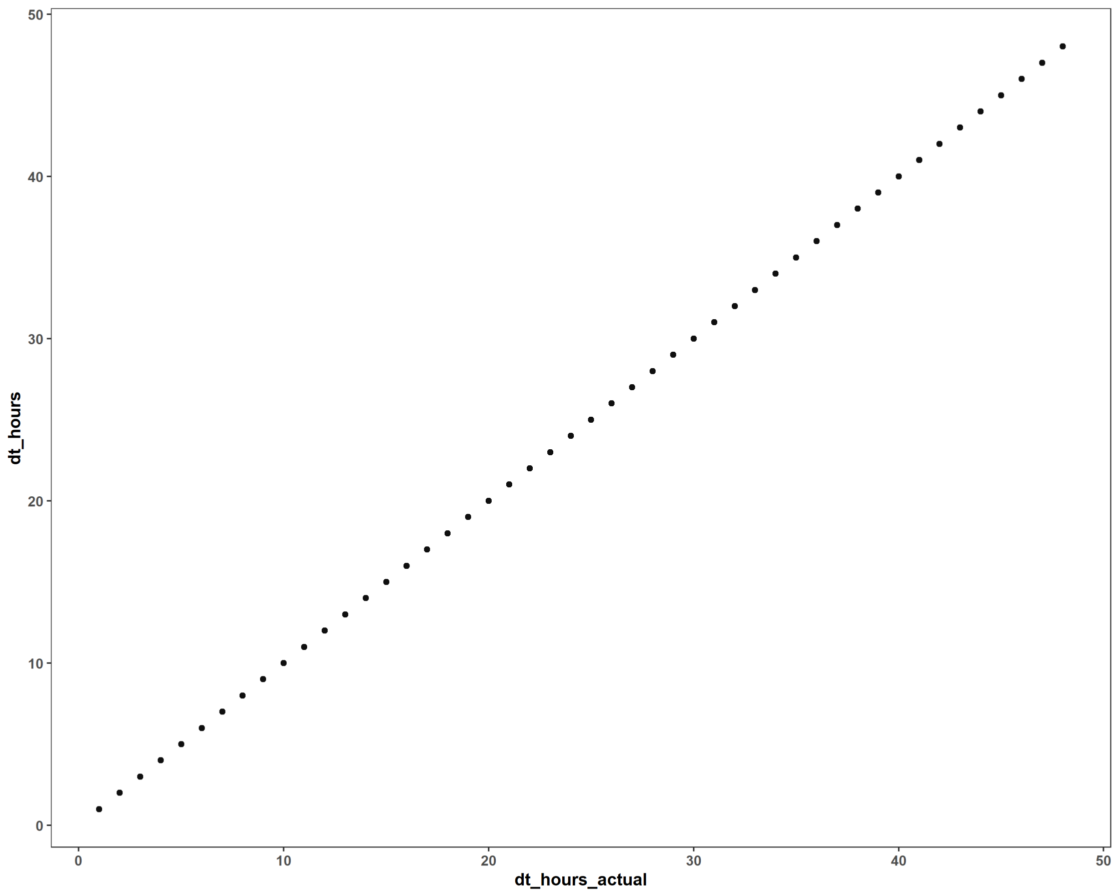


```r
# diagnostic check of speed vs diff with changing sample size
d %>%
  filter(is.finite(speed_est)) %>%
  ggplot(aes(diff_est, speed_est)) +
  geom_errorbar(aes(ymin = speed_lwr_95, ymax = speed_upr_95),
                alpha = 0.2, width = 0) +
  geom_errorbarh(aes(xmin = diff_lwr_95, xmax = diff_upr_95),
                 alpha = 0.2, height = 0) +
  geom_point(aes(color = dt_hours)) +
  geom_point(size = 2, shape = 1) +
  geom_smooth(aes(group = dt_hours, color = dt_hours), method = 'lm',
              formula = y ~ x, se = FALSE) +
  scale_x_continuous('Estimated diffusion (km\U00B2/day, log scale)',
                     transform = 'log', labels = \(x) round(x, 2)) +
  scale_y_continuous('Estimated speed (km/day, log scale)',
                     transform = 'log', labels = \(x) round(x, 2)) +
  khroma::scale_color_smoothrainbow(name = 'Sampling interval (hours)',
```

```
                                    breaks = c(1, 10, 20, 30, 40, 48),
                                    range = c(0.1, 1)) +
  theme(legend.position = 'top')
```

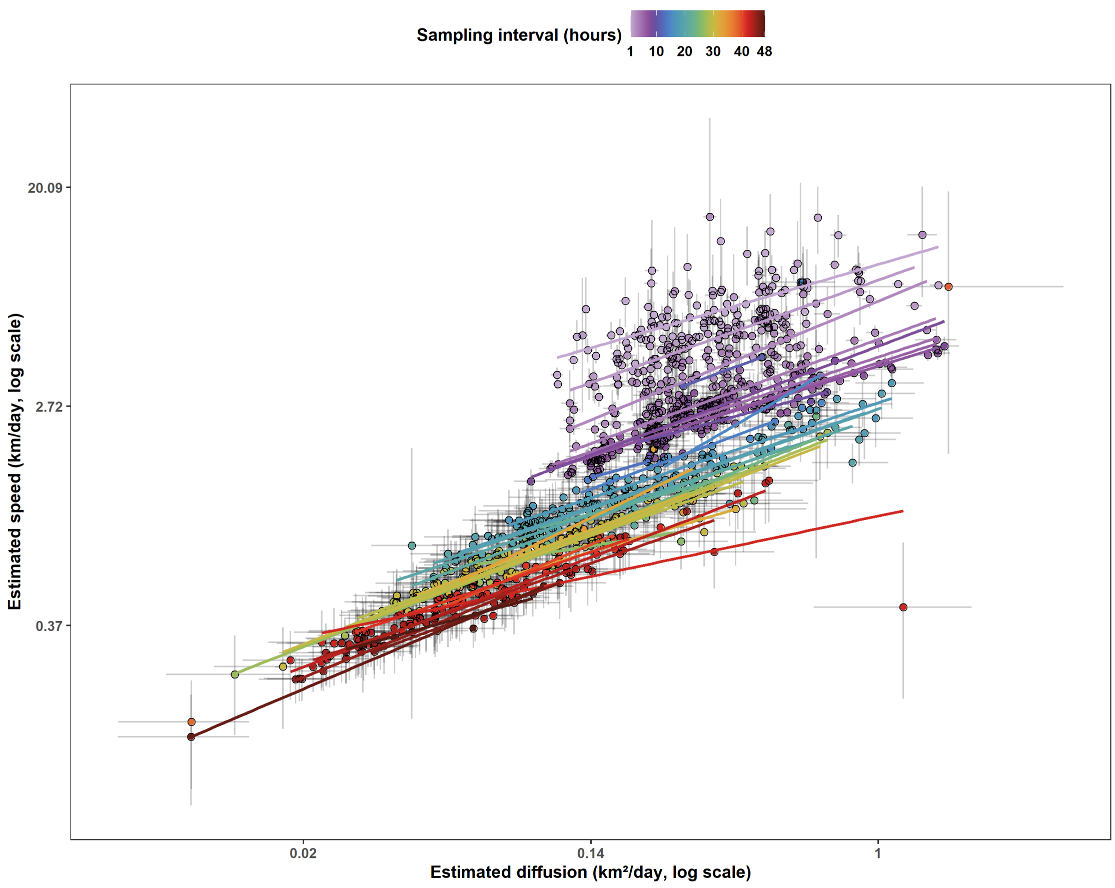


```
# count number of sampling intervals with < 40 estimates of 117 telemetries
n_distinct(d$animal_year)

## [1] 117

d %>%
  group_by(dt_hours) %>%
  summarise(n = sum(is.finite(speed_est))) %>%
  filter(n <= 40) %>%
  nrow()

## [1] 41

d %>%
  group_by(dt_hours) %>%
  summarise(n = sum(is.finite(diff_est))) %>%
  filter(n <= 40) %>%
  nrow()

## [1] 0
```

```r
# get directional persistence & range crossing times for full tels ----
d_0 <-
  readRDS('../ny-deer-vasectomy/models/full-telemetry-movement-models-2024-04
-20.rds') %>%
  mutate(animal_year = paste(animal, study.year)) %>%
  filter(! duplicated(animal_year)) %>% # had some accidental duplicates
  filter(animal_year %in% as.character(d$animal_year)) %>%
  mutate(tau_v = map_dbl(model, \(.m) {
    ci <- summary(.m, units = FALSE)$CI
    if(any(rownames(ci) == 'τ[velocity] (seconds)')) {
      tv <- ci['τ[velocity] (seconds)', 'est']
      tv <- 'h' %#% tv
    } else {
      tv <- NA_real_
    }
    return(tv)
  }),
  tau_p = map_dbl(model, \(.m) {
    ci <- summary(.m, units = FALSE)$CI
    if(any(rownames(ci) == 'τ[position] (seconds)')) {
      tp <- ci['τ[position] (seconds)', 'est']
      tp <- 'hours' %#% tp
    } else {
      tp <- NA_real_
    }
    return(tp)
  }))

# almost all deer have a tau_v < 1 hour or do not have a speed estimate
nrow(d_0)

## [1] 117

mean(d_0$tau_v, na.rm = TRUE) * 60

## [1] 10.21515

quantile(d_0$tau_v, na.rm = TRUE) * 60

##        0%       25%       50%       75%      100%
##  2.506970  7.485053  9.748160 12.617718 26.902792

hist(d_0$tau_v * 60, main = NULL, xlab = expression(tau[v]~(minutes)))
```

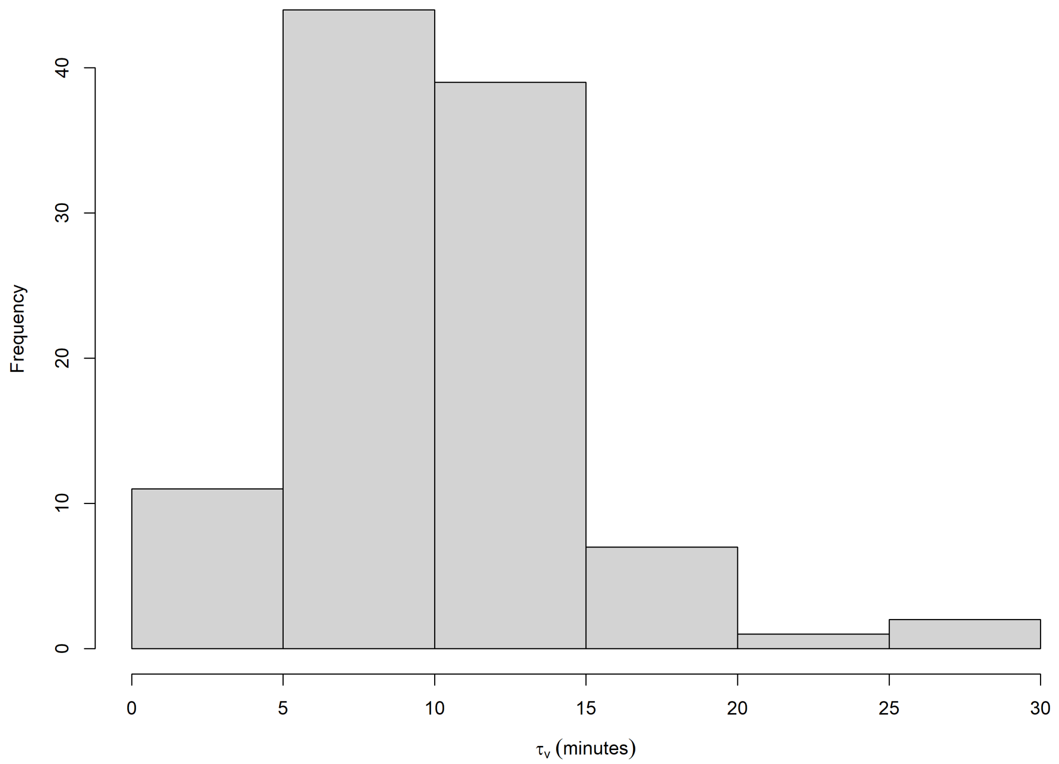


```r
all(d_0$tau_v < 1, na.rm = TRUE)
```

```
## [1] TRUE
```

```r
all(d_0$tau_v < 1 | is.na(d_0$tau_v))
```

```
## [1] TRUE
```

```r
# find stats for range crossing time in days
mean(d_0$tau_p, na.rm = TRUE)
```

```
## [1] 27.83583
```

```r
quantile(d_0$tau_p, na.rm = TRUE)
```

```
##         0%        25%        50%        75%       100%
##   3.692419   6.133823   9.128437  14.853723 797.963836
```

```r
max(d_0$tau_p, na.rm = TRUE) / 24
```

```
## [1] 33.24849
```

```r
hist(d_0$tau_p, main = NULL, xlab = expression(tau[v]~(minutes)))
```

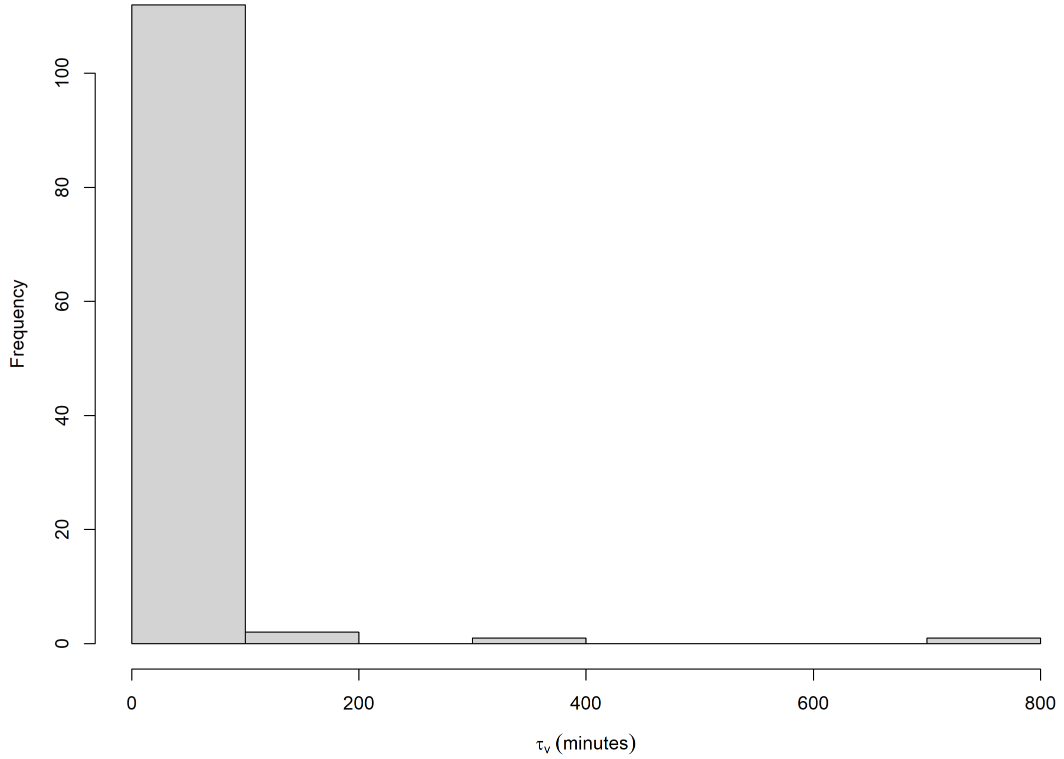


```
rm(d_0) # to save space

# fit GAM ----
# not using weights because they bias towards smaller tau values
m <- bam(speed_est ~
           log(diff_est) +
           s(dt_hours, k = 10, bs = 'tp') +
           s(log(diff_est), animal_year, bs = 'fs'),
         data = filter(d, is.finite(speed_est)),
         family = Gamma(link = 'log'),
         method = 'fREML',
         discrete = TRUE)
draw(m, overall_uncertainty = FALSE, parametric = TRUE)
```

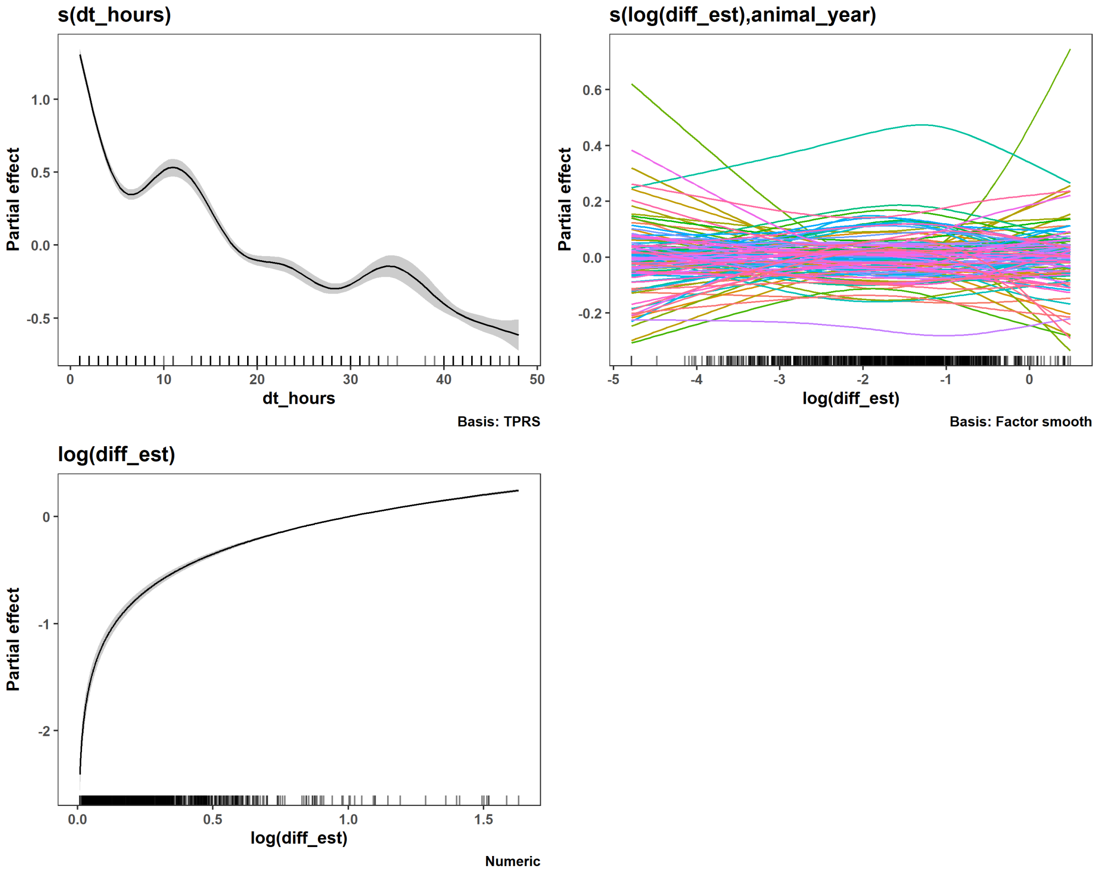


```r
appraise(m, point_alpha = 0.1, n_bins = 30)
```

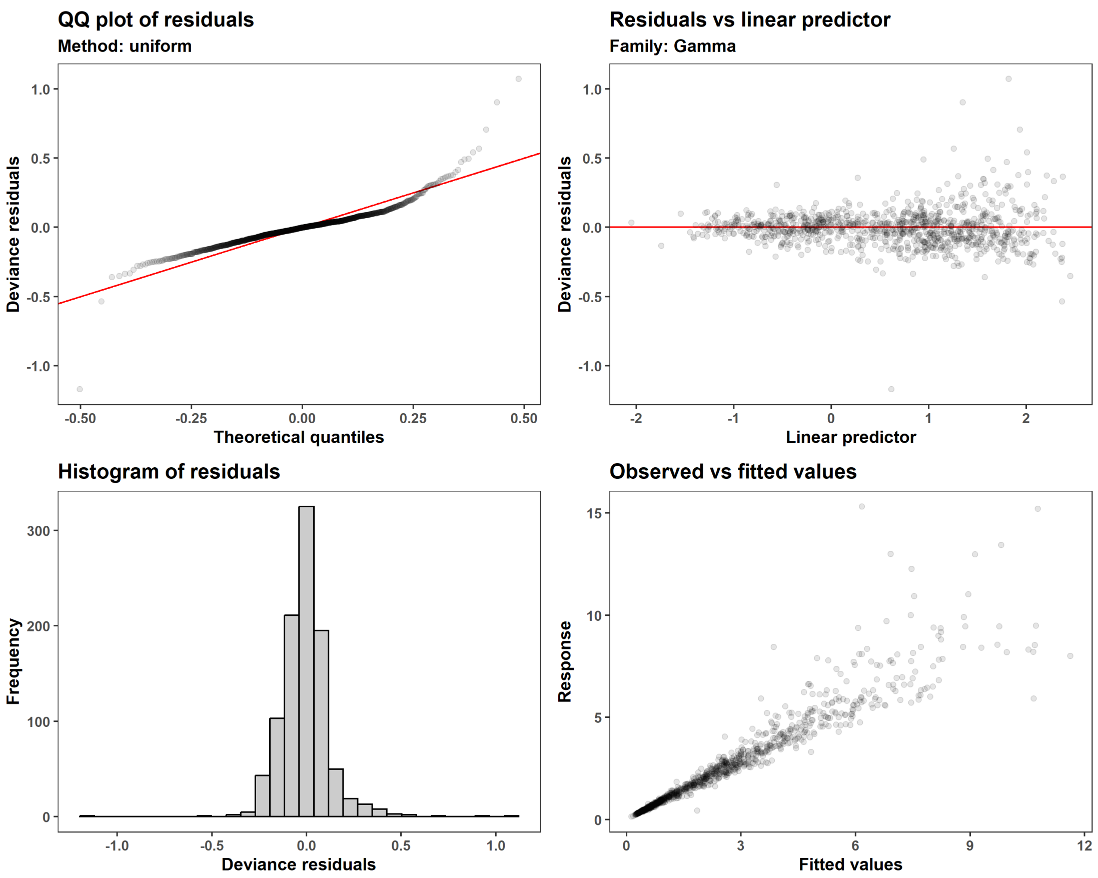


```
summary(m)

##
## Family: Gamma
## Link function: log
##
## Formula:
## speed_est ~ log(diff_est) + s(dt_hours, k = 10, bs = "tp") +
##     s(log(diff_est), animal_year, bs = "fs")
##
## Parametric coefficients:
##                Estimate Std. Error t value Pr(>|t|)
## (Intercept)     1.16406    0.03412   34.11   <2e-16 ***
## log(diff_est)  0.50407    0.01584   31.82   <2e-16 ***
## ---
## Signif. codes:  0 '***' 0.001 '**' 0.01 '*' 0.05 '.' 0.1 ' ' 1
##
## Approximate significance of smooth terms:
##                                  edf  Ref.df       F p-value
## s(dt_hours)                    8.844   8.982 595.547  <2e-16 ***
## s(log(diff_est),animal_year) 116.368 681.000   0.631  <2e-16 ***
## ---
## Signif. codes:  0 '***' 0.001 '**' 0.01 '*' 0.05 '.' 0.1 ' ' 1
```

```
##
## R-sq.(adj) =  0.888   Deviance explained = 97.7%
## fREML = -365.74  Scale est. = 0.022629  n = 984
```

```r
# est and CIs for slope of log(diffusion)
coef(m)['log(diff_est)']
```

```
## log(diff_est)
##     0.5040739
```

```r
coef(m)['log(diff_est)']+summary(m)$se['log(diff_est)']*qnorm(c(0.005,0.995))
```

```
## [1] 0.4632738 0.5448741
```

```r
# plot on log-log scale
p_speed_diff_2log <-
  expand_grid(animal_year = 'new animal',
              dt_hours = c(1, 10, 20, 30, 40, 48),
              diff_est = exp(seq(log(min(d$diff_est)), log(2.34),
                                 length.out = 400))) %>%
  bind_cols(.,
            predict(m, newdata = ., se.fit = TRUE, type = 'link',
                    unconditional = FALSE, discrete = FALSE,
                    exclude = 's(log(diff_est),animal_year)') %>%
              as.data.frame() %>%
              transmute(mu = exp(fit),
                        lwr = exp(fit + qnorm(0.005) * se.fit),
                        upr = exp(fit + qnorm(0.995) * se.fit))) %>%
  ggplot() +
  geom_point(aes(diff_est, speed_est, color = dt_hours), d, alpha = 0.3) +
  geom_ribbon(aes(diff_est, ymin = lwr, ymax = upr, fill = dt_hours,
                  color = dt_hours, group = dt_hours), alpha = 0.5, lwd=0.3)+
  geom_line(aes(diff_est, mu, group = dt_hours), lwd = 0.75) +
  geom_line(aes(diff_est, mu, color = dt_hours, group = dt_hours)) +
  scale_x_continuous('Estimated diffusion (km\U00B2/day, log scale)',
                     transform = 'log',labels = \(x) round(exp(x), 2)) +
  scale_y_continuous('Estimated speed (km/day, log scale)',
                     transform = 'log', labels = \(x) round(exp(x), 2)) +
  scale_color_iridescent(name = 'Sampling interval (hours)',
                         limits = c(1, 48), reverse = TRUE,
                         breaks = c(1, 10, 20, 30, 40, 48),
                         range = c(0.1, 0.6)) +
  scale_fill_iridescent(name = 'Sampling interval (hours)',
                        limits = c(1, 48), reverse = TRUE,
                        breaks = c(1, 10, 20, 30, 40, 48),
                        range = c(0.1, 0.6)) +
  theme(legend.position = 'top')
```

```
## Warning in predict.gam(object, newdata = newdata, type = type, se.fit = se
.fit,
## : factor levels new animal not in original fit
```

```r
# plot on the natural scale
p_speed_diff <-
  expand_grid(animal_year = 'new animal',
              dt_hours = c(1, 10, 20, 30, 40, 48),
              diff_est = seq(1e-10, 2.34, length.out = 400)) %>%
  bind_cols(.,
            predict(m, newdata = ., se.fit = TRUE, type = 'link',
                    unconditional = FALSE, discrete = FALSE,
                    exclude = 's(log(diff_est),animal_year)') %>%
              as.data.frame() %>%
              transmute(mu = exp(fit),
                        lwr = exp(fit + qnorm(0.005) * se.fit),
                        upr = exp(fit + qnorm(0.995) * se.fit))) %>%
  ggplot() +
  geom_point(aes(diff_est, speed_est, color = dt_hours), d, alpha = 0.3) +
  geom_ribbon(aes(diff_est, ymin = lwr, ymax = upr, fill = dt_hours,
                  color = dt_hours, group = dt_hours), alpha = 0.5, lwd=0.3)+
  geom_line(aes(diff_est, mu, group = dt_hours), lwd = 1) +
  geom_line(aes(diff_est, mu, color = dt_hours, group = dt_hours)) +
  scale_x_continuous('Estimated diffusion (km\U00B2/day)') +
  scale_y_continuous('Estimated speed (km/day)') +
  scale_color_iridescent(name = 'Sampling interval (hours)',
                         limits = c(1, 48), reverse = TRUE,
                         breaks = c(1, 10, 20, 30, 40, 48),
                         range = c(0.1, 0.6)) +
  scale_fill_iridescent(name = 'Sampling interval (hours)',
                        limits = c(1, 48), reverse = TRUE,
                        breaks = c(1, 10, 20, 30, 40, 48),
                        range = c(0.1, 0.6)) +
  theme(legend.position = 'inside', legend.position.inside = c(0.5, 0.95),
        legend.direction = 'horizontal')

## Warning in predict.gam(object, newdata = newdata, type = type, se.fit = se
.fit,
## : factor levels new animal not in original fit

# estimated parameters vs sampling interval ----
# speed vs sampling interval
p_speed_dt <-
  d %>%
  ggplot(aes(dt_hours, speed_est)) +
  geom_jitter(height = 0, width = 0.25, size = 0.3, alpha = 0.2) +
  geom_smooth(method = 'gam', formula = y ~ s(x, bs = 'ts', k = 5),
              method.args = list(family = Gamma(link = 'log')),
              color = '#DDAA33', fill = '#DDAA33', alpha = 0.3,
              level = 0.99) +
  scale_x_continuous('Sampling interval (hours)', limits = c(0, 49)) +
  scale_y_continuous('Estimated speed (km/day)')

# diff vs sampling interval
```

```r
p_diff_dt <-
  d %>%
  ggplot(aes(dt_hours, diff_est)) +
  geom_jitter(height = 0, width = 0.25, size = 0.3, alpha = 0.2) +
  geom_smooth(method = 'gam', formula = y ~ s(x, bs = 'ts', k = 5),
              method.args = list(family = Gamma(link = 'log')),
              color = '#DDAA33', fill = '#DDAA33', alpha = 0.3,
              level = 0.99) +
  scale_x_continuous('Sampling interval (hours)', limits = c(0, 49)) +
  scale_y_continuous('Estimated diffusion (km\U00B2/day)')

# degrees of freedom vs sampling interval ----
p_speed_dof_dt <-
  (d %>%
     filter(is.finite(speed_est)) %>%
     ggplot(aes(dt_hours, speed_dof / duration)) +
     coord_cartesian(ylim = c(0, 20)) +
     geom_jitter(height = 0, width = 0.25, size = 0.3, alpha = 0.2) +
     geom_smooth(method = 'gam', formula = y ~ s(x, bs = 'ad', k = 10),
                 method.args = list(family = Gamma(link = 'log')),
                 color = '#DDAA33', fill = '#DDAA33', alpha = 0.3,
                 level = 0.99) +
     scale_x_continuous('Sampling interval (hours)', limits = c(0, 49)) +
     scale_y_continuous('Speed ESS per day')) %>%
  ggMarginal(margins = 'x', fill = 'grey', type = 'histogram',
             binwidth = 1, center = 1)

p_diff_dof_dt <-
  (d %>%
     filter(is.finite(diff_est)) %>%
     ggplot(aes(dt_hours, diff_dof / duration)) +
     coord_cartesian(ylim = c(0, 20)) +
     geom_jitter(height = 0, width = 0.25, size = 0.3, alpha = 0.2) +
     geom_smooth(method = 'gam', formula = y ~ s(x, bs = 'ad', k = 10),
                 method.args = list(family = Gamma(link = 'log')),
                 color = '#DDAA33', fill = '#DDAA33', alpha = 0.3,
                 level = 0.99) +
     scale_x_continuous('Sampling interval (hours)', limits = c(0, 49)) +
     scale_y_continuous('Diffusion ESS per day')) %>%
  ggMarginal(margins = 'x', fill = 'grey', type = 'histogram',
             binwidth = 1, center = 1)

# plot everything together
fig_3 <-
  plot_grid(p_speed_diff,
            plot_grid(p_speed_dt, p_speed_dof_dt, p_diff_dt, p_diff_dof_dt,
                      ncol = 2, labels = LETTERS[2:5],
                      rel_heights = c(1.2, 1)),
            labels = c('A', ''), nrow = 1)
```

```
## Warning: Removed 4632 rows containing non-finite outside the scale range
## (`stat_smooth()`).

## Warning: Removed 1397 rows containing non-finite outside the scale range
## (`stat_smooth()`).

plot(1)
```

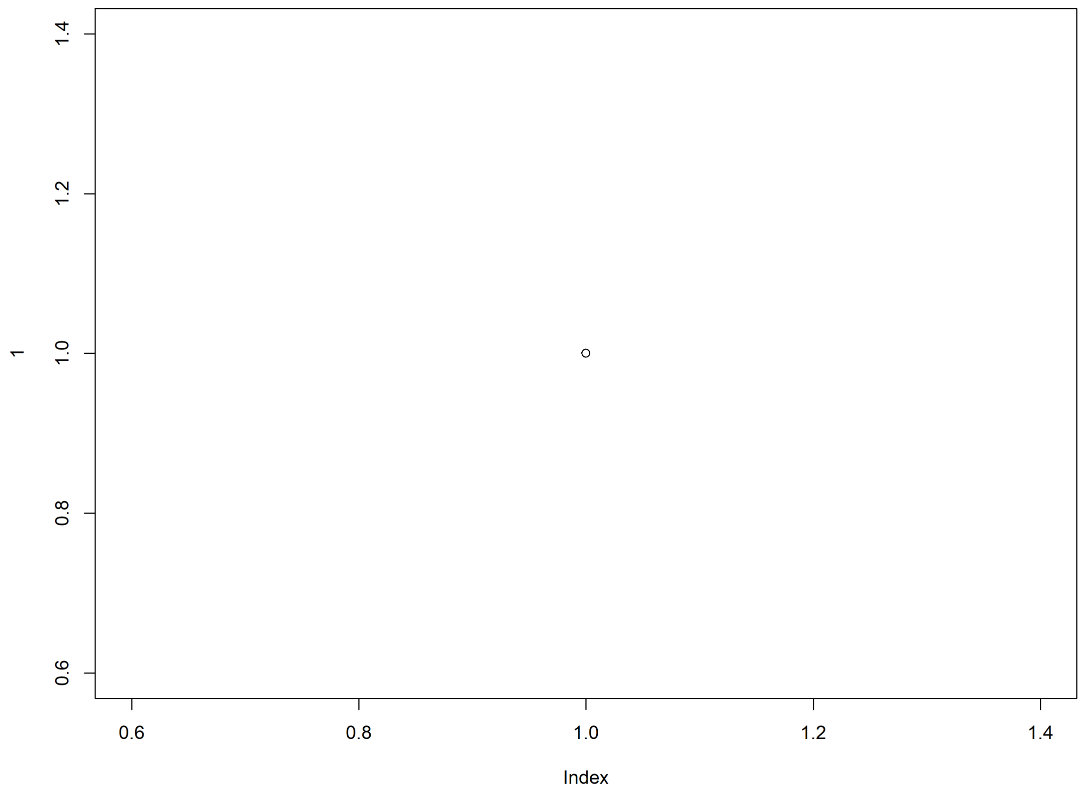


```
fig_3
```

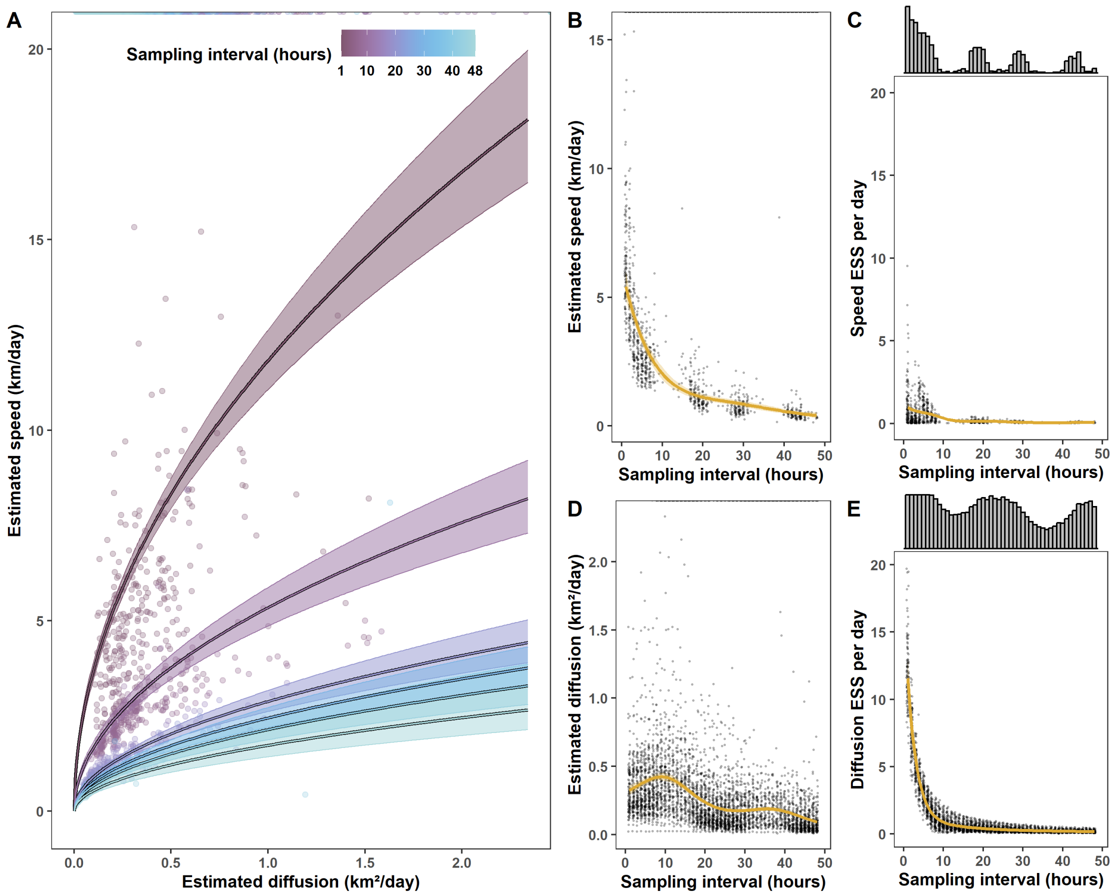


```r
# find percentage of (n_eff / day) > 5
mean(d$speed_dof / d$duration > 5) * 100

## [1] 0.08903134

mean(d$diff_dof / d$duration > 5) * 100

## [1] 5.48433

all(filter(d, dt_hours == 1)$diff_dof / filter(d, dt_hours == 1)$duration>5)

## [1] TRUE
```

### 5 Simulations (Fig.3)

```r
source('analysis/figures/default-ggplot-theme.R') # gets reset in new chunk
tau_p <- 24 %#% 'hour'    # true range crossing time
tau_v <- 1 %#% 'hour'    # true directional persistence
delta_t_0 <- 1 %#% 'hour' # finest sampling time
t_start <- as.POSIXct('2024-01-01 00:00')
TIMES <- seq(t_start, t_start + 100 %#% 'days', by = delta_t_0)
range(TIMES)

## [1] "2024-01-01 00:00:00 PST" "2024-04-10 01:00:00 PDT"

diff(range(TIMES))

## Time difference of 100 days

N_DAYS <- as.numeric(diff(range(TIMES)))

m_0 <- ctmm(tau = c(tau_p, tau_v),
            mu = c(0, 0), # centered at (0, 0)
            sigma = 1e6) # variance for the bivariate gaussian HR

# extract the true speed and diffusion values
summary(m_0)

## $name
## [1] "OUF anisotropic"
##
## $DOF
##      mean      area diffusion     speed
##         0         0         0         0
##
## $CI
##                                    low       est high
## area (square kilometers)             0 18.822741  Inf
## τ[position] (hours)                  0 24.000000  Inf
## τ[velocity] (hours)                  0  1.000000  Inf
## speed (kilometers/day)               0  6.928203  Inf
## diffusion (square kilometers/day)    0  1.741890  Inf

speed_0 <- 'km/day' %#% speed(m_0, units = FALSE)$CI[1, 'est']
diff_0 <- 'km^2/day' %#%
  summary(m_0, units = FALSE)$CI['diffusion (square meters/second)', 'est']

# set number of simulations to run
a <- 0.03
(1/(a))^2 #' find number of sims necessary to be within `a * 100%` of truth

## [1] 1111.111

n_sims <- 1e3
paste(round(1/sqrt(n_sims) * 100, 2), '%') # find approximate accuracy
```

```r
## [1] "3.16 %"

# set number of cores to use
NCORES <- min(12, availableCores(logical = FALSE) - 2)
plan(multisession, workers = NCORES)

# set file names
sims_name <- paste0('data/simulations/1e', log10(n_sims),
                    '-simulated-telemetries.rds')
models_name <- paste0('models/simulations/1e', log10(n_sims),
                      '-movement-models.rds')

if(file.exists(models_name)) {
  sims <- readRDS(models_name) # import models
} else {
  if(file.exists(sims_name)) {
    tels <- readRDS(sims_name) # import simulated telemetries w/o models
  } else {
    tels <- tibble(seed = 1:n_sims,
                   tel = map(seed, \(.seed) {
                     library('ctmm')
                     simulate(m_0, nsim = 1, seed = .seed, # see seed below
                              t = as.numeric(TIMES)) %>%
                       return()
                   }, .progress = TRUE))

    layout(matrix(1:4, ncol = 2))
    plot(tels$tel[[1]], error = FALSE, type = 'l') # bivariate SD is 1 km
    plot(tels$tel[[2]], error = FALSE, type = 'l')
    plot(tels$tel[[3]], error = FALSE, type = 'l')
    plot(tels$tel[[4]], error = FALSE, type = 'l')
    layout(1)
    saveRDS(tels, sims_name)
  }

  # fit movement models and extract estimates
  #' note: `future_map_*()` slows down fast computations
  map_int(48:1, \(.i) {
    tels %>%
      mutate(
        delta_t = .i, # add column of sampling time
        tel = map2(tel, delta_t, \(.tel, .dt) {
          if(.dt == 1) {
            return(.tel)
          } else {
            # starting from mod == 1 to ensure max number of samples
            return(.tel[1:length(TIMES) %% .dt == 1, ])
          }
        }, .progress = 'Thinning'),
```

```r
        n_fixes = map_int(tel, nrow),
        vg = map(tel, \(x) ctmm.guess(data = x, interactive = FALSE),
                 .progress = 'Variograms'),
        mm = future_map2(tel, vg, \(.t, .v) {
          #' using `ctmm.fit()` rather than `ctmm.select()` to avoid issues
          #' due to missingness *not* at random of fast realizations
          #' this will cause CIs to sometimes include `Inf` and be
          #' positively biased, on average
          #' using explicit `return()` to avoid having a warning returned
          return(ctmm.fit(data = .t, CTMM = .v))
        }, .progress = TRUE, .options = furrr_options(seed = NULL)),
        estimates = map(mm, \(.m) {
          # extract speed estimates
          .speed <-
            speed(.m, units = FALSE)$CI %>%
            as.data.frame() %>%
            transmute(speed_lwr = 'km/day' %#% low,
                      speed_est = 'km/day' %#% est,
                      speed_upr = 'km/day' %#% high) %>%
            suppressWarnings() # to avoid warnings on fractal movement

          cis <- as.data.frame(summary(.m, units = FALSE)$CI)

          # extract root-mean-squared speed estimates
          # (summary sometimes doesn't have speed in ctmm version 1.2.1)
          if(any(rownames(cis) == 'speed (meters/second)')) {
            .rms_speed <-
              slice(cis, which(rownames(cis) == 'speed (meters/second)')) %>%
              transmute(rms_speed_lwr = 'km/day' %#% low,
                        rms_speed_est = 'km/day' %#% est,
                        rms_speed_upr = 'km/day' %#% high)
          } else {
            .rms_speed <-
              tibble(rms_speed_lwr = NA_real_,
                     rms_speed_est = NA_real_,
                     rms_speed_upr = NA_real_)
          }

          # extract diffusion estimates
          .diff <-
            slice(cis,
                  which(rownames(cis)=='diffusion (square meters/second)'))%>
%
            transmute(diff_lwr = 'km^2/day' %#% low,
                      diff_est = 'km^2/day' %#% est,
                      diff_upr = 'km^2/day' %#% high)

          # rownames are unnecessary
          rownames(.speed) <- rownames(.rms_speed) <- rownames(.diff) <- NULL
```

```
        return(bind_cols(.speed, .rms_speed, .diff))
      })) %>%
    unnest(estimates) %>%
    mutate(speed_lwr = if_else(speed_lwr == 0, NA_real_, speed_lwr),
           speed_est = if_else(is.infinite(speed_est), NA_real_, speed_est)
,
           speed_upr = if_else(is.infinite(speed_upr), NA_real_,
                               speed_upr)) %>%
    saveRDS(gsub('.rds', paste0('-', .i, '-h.rds'), models_name))

    return(.i)
  })

  # save all the movement models to a single data frame
  sims <- map_dfr(1:48, \(.i) {
    gsub('.rds', paste0('-', .i, '-h.rds'), models_name) %>%
      readRDS()
  }) %>%
    #' change `NA`s to `0` and `Inf`
    mutate(speed_lwr = if_else(is.na(speed_lwr), 0, speed_lwr),
           speed_est = if_else(is.na(speed_est), Inf, speed_est),
           speed_upr = if_else(is.na(speed_upr), Inf, speed_upr),
           diff_lwr = if_else(is.na(diff_lwr), 0, diff_lwr),
           diff_est = if_else(is.na(diff_est), Inf, diff_est),
           diff_upr = if_else(is.na(diff_upr), Inf, diff_upr)) %>%
    mutate(
      n_eff_speed = future_map_dbl(mm, \(.m) summary(.m)$DOF['speed'],
                                   .progress = TRUE),
      n_eff_diff = future_map_dbl(mm, \(.m) summary(.m)$DOF['diffusion'],
                                  .progress = TRUE))

  saveRDS(sims, models_name)

  Sys.time()
}

# find summary statistics for each dt
sims_sum <- sims %>%
  group_by(delta_t) %>%
  summarize(n_fixes = unique(n_fixes),
            n_speed = sum(! is.na(speed_est)),
            n_diff = sum(! is.na(diff_est)),
            speed_lwr_99 = quantile(speed_est, 0.005, na.rm = TRUE),
            speed_mean = mean(speed_est, na.rm = TRUE),
            speed_median = median(speed_est, na.rm = TRUE),
            speed_upr_99 = quantile(speed_est, 0.995, na.rm = TRUE),
            diff_lwr_99 = quantile(diff_est, 0.005, na.rm = TRUE),
            diff_mean = mean(diff_est, na.rm = TRUE),
```

```r
            diff_median = median(diff_est, na.rm = TRUE),
            diff_upr_99 = quantile(diff_est, 0.995, na.rm = TRUE),
            n_eff_speed_lwr_99 = quantile(n_eff_speed, 0.005, na.rm = TRUE),
            n_eff_speed = mean(n_eff_speed),
            n_eff_speed_upr_99 = quantile(n_eff_speed, 0.995, na.rm = TRUE),
            n_eff_diff_lwr_99 = quantile(n_eff_diff, 0.005, na.rm = TRUE),
            n_eff_diff = mean(n_eff_diff),
            n_eff_diff_upr_99 = quantile(n_eff_diff, 0.995, na.rm = TRUE))

# create figures of the results
tau_p_h <- 'hours' %#% tau_p
K <- 3 #' `ylim = c(0, truth * K)`

if(FALSE) {
  # check relationship as delta_t changes
  library(mgcv)
  gam_0 <- gam(speed_est ~ log(diff_est),
               family = Gamma(link = 'log'),
               data = sims,
               subset = delta_t == 1,
               method = 'REML')
  summary(gam_0)

  max_dt <- 5

  sims %>%
    filter(delta_t <= max_dt) %>%
    summarise(nas = sum(is.na(c(diff_est, speed_est))),
              infs = sum(is.infinite(c(diff_est, speed_est))))

  #' relationship between speed and diffusion deteriorates if we use
  #' `ctmm.fit()` instead of `ctmm.select()`
  sims %>%
    filter(delta_t <= max_dt) %>%
    mutate(delta_t = paste0('\U0394', 't = ', delta_t, ' h')) %>%
    ggplot(aes(diff_est, speed_est)) +
    facet_wrap(~ delta_t, scales = 'free') +
    geom_point(alpha = 0.1) +
    geom_smooth(alpha = 0.1, n = 400) +
    labs(x = expression(bold(Estimated~diffusion~(m^2/s))),
         y = expression(bold(Estimated~speed~(m/s))))
}

p_a <-
  ggplot(sims_sum, aes(delta_t)) +
  coord_cartesian(ylim = c(0, K * speed_0)) +
  # geom_vline(xintercept = tau_p_h, lty = 'dashed', alpha = 0.5) +
  geom_errorbar(aes(ymin = speed_lwr_99, ymax = speed_upr_99),
                width = 0.5, alpha = 0.3) +
```

```r
  geom_hline(yintercept = speed_0, color = 'red3') +
  geom_line(aes(y = speed_median), alpha = 0.2) +
  geom_point(aes(y = speed_median)) +
  labs(x = 'Sampling interval (hours)',
       y = 'Estimated speed (km/day)')

p_b <-
  ggplot(sims_sum, aes(delta_t, diff_mean)) +
  coord_cartesian(ylim = c(0, K * diff_0)) +
  # geom_vline(xintercept = tau_p_h, lty = 'dashed', alpha = 0.5) +
  geom_errorbar(aes(ymin = diff_lwr_99, ymax = diff_upr_99),
                width = 0.5, alpha = 0.3) +
  geom_hline(yintercept = diff_0, color = 'red3') +
  geom_line(aes(y = diff_median), alpha = 0.2) +
  geom_point(aes(y = diff_median)) +
  labs(x = 'Sampling interval (hours)',
       y = 'Estimated diffusion (km\U00B2/day)')

#' force negative ESS to 0. the issue is now corrected in ctmm, but the
#' negative values are so few
mean(sims$n_eff_diff < 0)

## [1] 0.0002708333

mean(sims$n_eff_speed < 0)

## [1] 2.083333e-05

sims <- mutate(sims,
               n_eff_diff = if_else(n_eff_diff < 0, 0, n_eff_diff),
               n_eff_speed = if_else(n_eff_speed < 0, 0, n_eff_speed))

p_eff_a <-
  (sims %>%
     filter(is.finite(speed_est)) %>%
     ggplot(aes(delta_t, n_eff_speed / N_DAYS)) +
     geom_jitter(height = 0, width = 0.25, size = 0.3, alpha = 0.05) +
     geom_smooth(method = 'gam', formula = y ~ s(x, bs = 'ad', k = 30),
                 # gamma family fails due to non-finite derivatives
                 # method.args = list(family = Gamma(link = 'log')),
                 color = '#DDAA33', fill = '#DDAA33', alpha = 0.3,
                 level = 0.99, data = filter(sims, n_eff_speed > 0),
                 n = 400, lwd = 1) +
     scale_x_continuous('Sampling interval (hours)', limits = c(0, 49)) +
     scale_y_continuous('Speed ESS per day',
                        breaks = seq(0, 24, by = 6), limits = c(0, 24))) %>%
  ggMarginal(margins = 'x', fill = 'grey', type = 'histogram',
             binwidth = 1, center = 1); p_eff_a

#' diffusion `n_eff` may jump as `delta_t` approaches the true diffusion
```

```r
p_eff_b <-
  (sims %>%
     filter(is.finite(diff_est)) %>%
     ggplot(aes(delta_t, n_eff_diff / N_DAYS)) +
     geom_jitter(height = 0, width = 0.25, size = 0.3, alpha = 0.05) +
     geom_smooth(method = 'gam', formula = y ~ s(x, bs = 'ad', k = 30),
                 # gamma family fails due to non-finite derivatives
                 # method.args = list(family = Gamma(link = 'log')),
                 color = '#DDAA33', fill = '#DDAA33', alpha = 0.3,
                 level = 0.99, data = filter(sims, n_eff_diff > 0),
                 n = 400, lwd = 1) +
     scale_x_continuous('Sampling interval (hours)', limits = c(0, 49)) +
     scale_y_continuous('Diffusion ESS per day',
                        breaks = seq(0, 24, by = 6), limits = c(0, 24))) %>%
  ggMarginal(margins = 'x', fill = 'grey', type = 'histogram',
             binwidth = 1, center = 1); p_eff_b
```

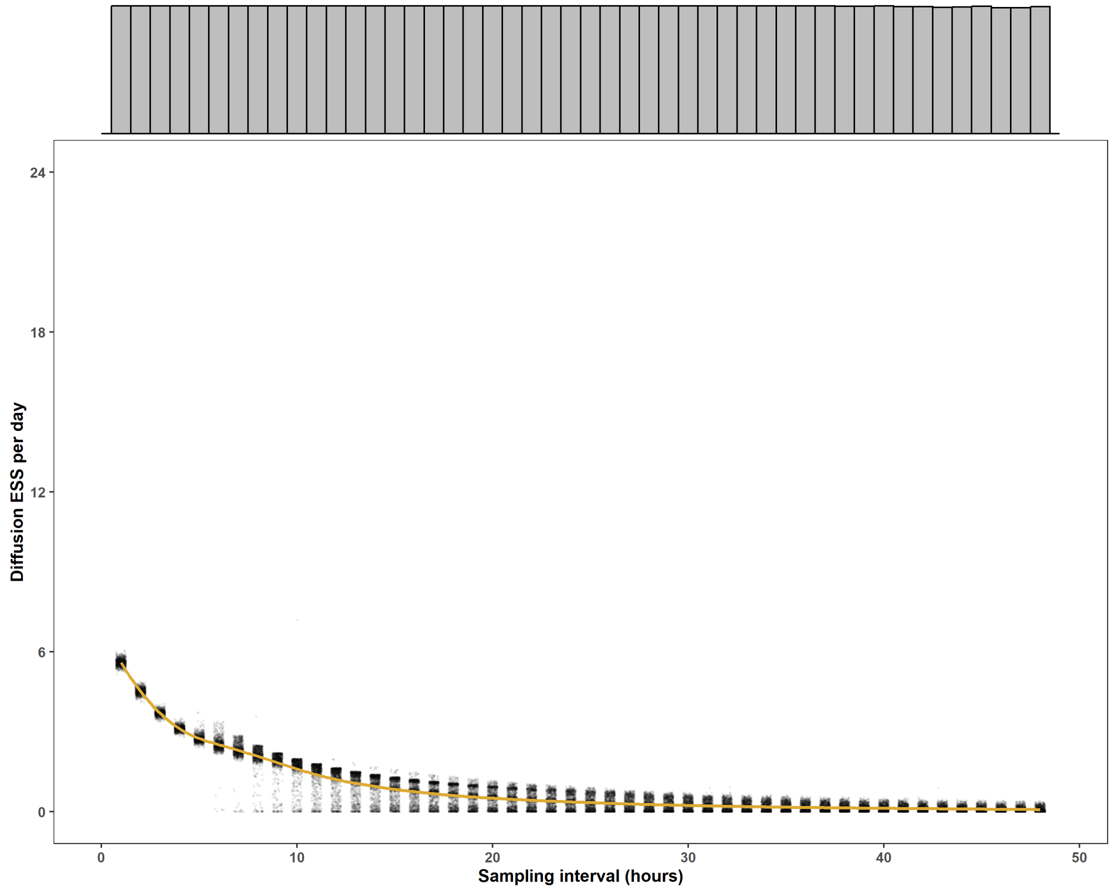


```r
plot_grid(p_a, p_eff_a, p_b, p_eff_b, ncol = 2, labels = 'AUTO')
```

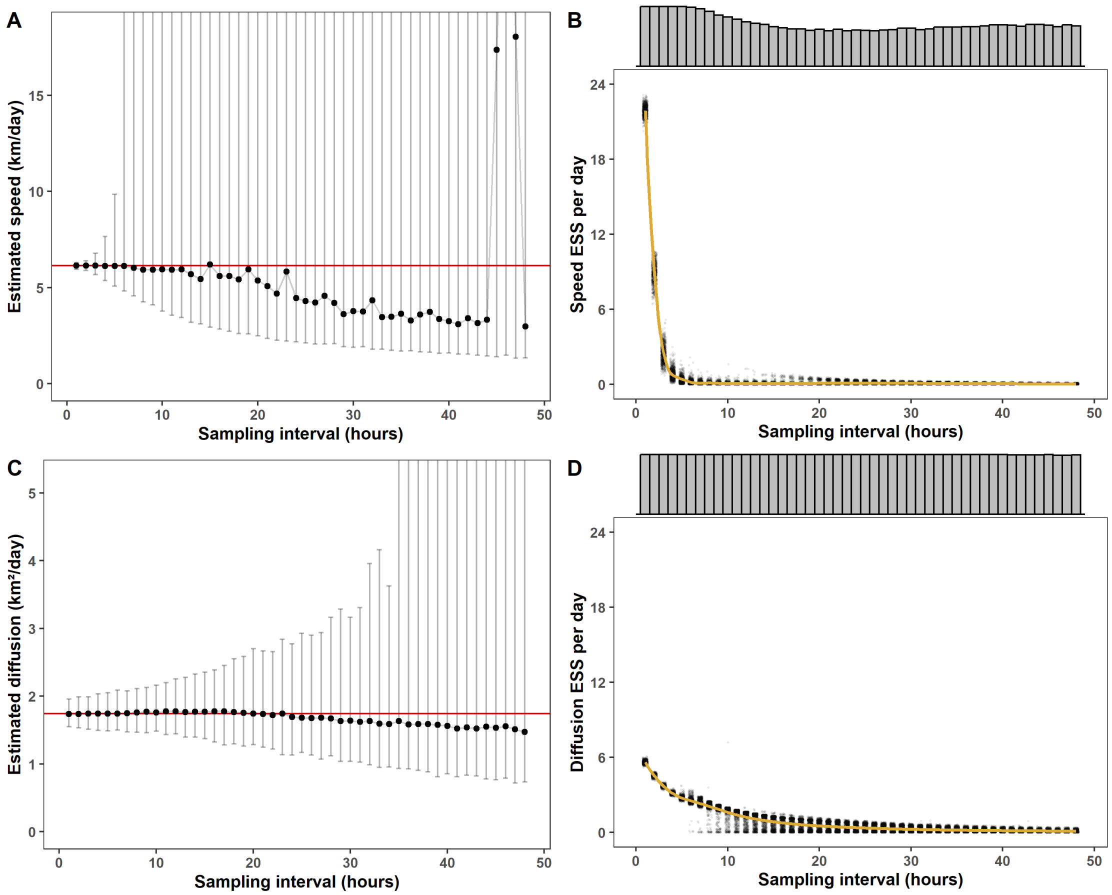


```
ggsave('figures/fig-3-diffusion-speed-sims.png', width = 12, height = 9,
       dpi = 600, bg = 'white')

ggplot() +
  geom_histogram(aes(delta_t, fill = 'all data'), alpha = 0.4,
                 color = 'black', sims, binwidth = 1) +
  geom_histogram(aes(delta_t, fill = 'finite diffusion'), alpha = 0.4,
                 color = 'black', filter(sims, is.finite(diff_est)),
                 binwidth = 1) +
  geom_histogram(aes(delta_t, fill = 'finite speed'), alpha = 0.4,
                 color = 'black', filter(sims, is.finite(speed_est)),
                 binwidth = 1) +
  scale_fill_brewer(type = 'qual', palette = 6)
```

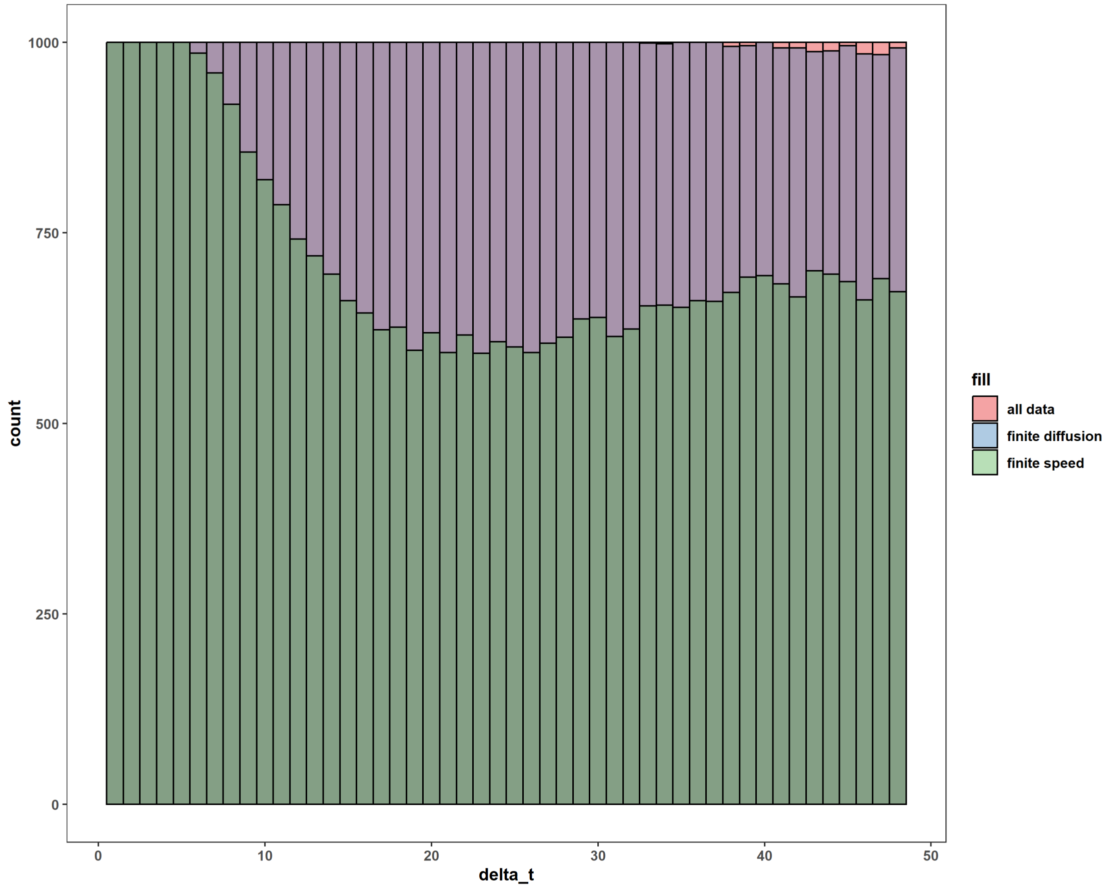


```
# find lowest dt values with infinite speed
min(filter(sims, is.infinite(speed_est))$delta_t)

## [1] 6

min(filter(sims, is.infinite(diff_est))$delta_t)

## [1] 33

sims %>%
  filter(is.finite(speed_est)) %>%
  ggplot(aes(delta_t)) +
  geom_histogram(binwidth = 1, center = 1)
```

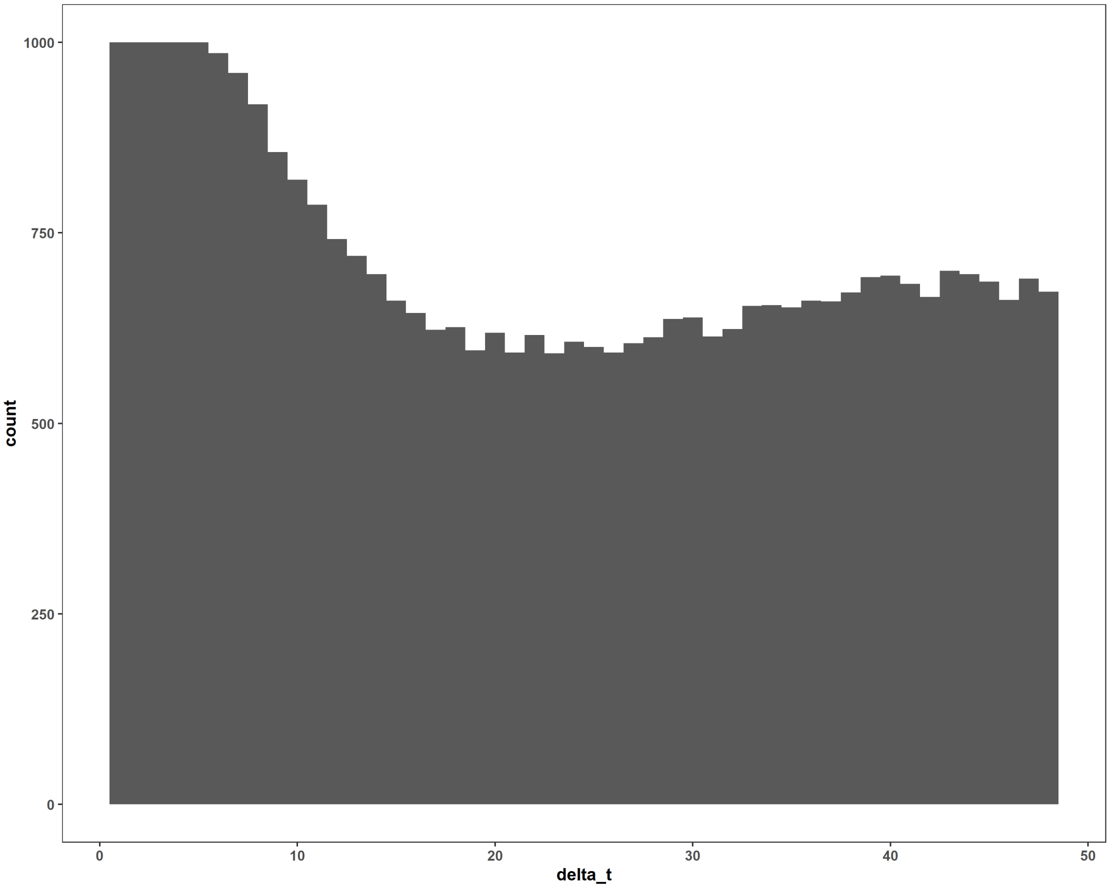

count
1000
750
500
250
0
0
10
20
30
40
50
delta_t
