## Supplement C - Interpreting the relationship between speed and diffusion for "Are your data too coarse for speed estimation? Diffusion rates as an alternative measure of animal movement"

### 1 Interpreting the relationship between speed and diffusion

The model we are using estimates log mean speed as a function of log diffusion:

$$\log(\mu) = \beta_0 + \beta_1 \log(d) \Longrightarrow \mu = \exp\big(\beta_0 + \beta_1 \log(d)\big) \qquad (1)$$

If we used a linear model of the form $\mu = \beta_0 + \beta_1 d$, the additive change in speed, $\Delta\mu$, for a change in diffusion, $\Delta d$, would be $\Delta\mu = \mu_1 - \mu_0 = (\beta_0 + (d + \Delta d)\beta_1) - (\beta_0 - d\beta_1) = (d + \Delta d - d)\beta_1 = \Delta d \beta_1$. In this case, $\beta_1$ indicates the additive change in $\mu$ for an additive change in diffusion, $\Delta d$.

If we used a GLM with a log link function, the model would be $\log(\mu) = \beta_0 + \beta_1 d \Longrightarrow \mu = \exp(\beta_0 + \beta_1 d)$, and the relative change in speed, $R_s$, for an additive change in diffusion, $\Delta d$ becomes

$$R_s = \mu_1/\mu_0 = \frac{\exp(\beta_0 + (d + \Delta d)d\beta_1)}{\exp(\beta_0 + d\beta_1)} = \frac{e^{\beta_0} e^{d\beta_1} e^{\Delta d \beta_1}}{e^{\beta_0} e^{d\beta_1}} = e^{\Delta d \beta_1}.$$

Now a $\Delta d$ additive change in diffusion would result in a relative change of $R_s = e^{\Delta d \beta_1}$. But since we log both speed and diffusion in our model, the relative change in speed is

$$R_s = \mu_1/\mu_0 = \frac{\exp\big(\beta_0 + \beta_1 \log(d + \Delta d)\big)}{\exp\big(\beta_0 + \beta_1 \log(d)\big)} = \exp\big(\beta_0 + \beta_1 \log(d + \Delta d) - \beta_0 - \beta_1 \log(d)\big) =$$

$$= \exp\left(\beta_1 \log\left(\frac{d + \Delta d}{d}\right)\right) = \exp\big(\beta_1 \log(R_d)\big) = \exp\left(\log\left(R_d^{\beta_1}\right)\right) = R_d^{\beta_1},$$

where $R_d$ is the relative change in diffusion and $\beta_1$ is the slope of the log-log model shown in Equation (1). Given the estimate from Fig. 1 of the manuscript $\hat{\beta}_1 = 0.40$, a 1% increase in diffusion corresponds to an approximate 1.004-fold increase (a 0.4% additive increase) in speed:

$$R_s = (101\%)^{0.40}/(100\%)^{0.40} = 1.01^{0.40}/1 \approx 1.003988 = 100.3988\% \approx 100.40\%.$$

This corresponds to an additive change in mean speed of $\Delta\mu \approx 100.40\% - 100\% = 0.40\%$, which is approximately $\beta_1$, but note the relationship between $\mu$ and d is nonlinear, so a doubling (a 100% increase) in diffusion does not correspond to a doubling in speed but rather an approximately 1.32-fold change (a 32% additive increase) in speed:

$$R_s = (200\%)^{0.40} = (2)^{0.40} = 1.319508 = 131.9508\% \Longrightarrow \Delta\mu = 31.9508\%.$$
